# The tumour suppressor RBM5 activates the helicase DHX15 to regulate splicing

**DOI:** 10.64898/2026.03.26.714623

**Authors:** Shiheng Liu, Tiantian Su, Jeffrey Huang, Chia-Ho Lin, Douglas L Black, Andrey Damianov, Z Hong Zhou

## Abstract

Pre-mRNA splicing determines the expressed proteome and is frequently dysregulated in cancer. The tumour-suppressor RBM5 controls an exon network regulating apoptosis, yet its molecular mechanism is elusive. Using in vivo spliceosome capture and cryogenic electron microscopy, we determined structures of precatalytic spliceosomes arrested by RBM5 immediately after U2 snRNP branchpoint recognition. Despite intron diversity, the U2–pre-mRNA duplex, branchpoint adenine, and downstream polypyrimidine tract are well-resolved. RBM5 binds the outer SF3B1 HEAT surface and performs dual functions: First, its helix–loop–helix motif and upstream zinc-finger domain sterically block tri-snRNP and Prp8 docking and prevent progression to pre-B and B^act^ complexes; Second, its G-patch activates DHX15 and places this DExH-box helicase on the pre-mRNA as it exits SF3B1, poised for branch helix unwinding. DHX15 binding to SF3B1 is facilitated by U2SURP/SR140, which engages SF3B1 near RBM5’s helix–loop–helix. Functional assays confirm that disruption of the RBM5 interfaces with either DHX15 or SF3B1 inhibit exon repression. Mutations at these regulatory interfaces are common in cancer genomes and predicted to disrupt its regulation of apoptotic isoforms. Thus, RBM5 acts as a dual-action spliceosome gatekeeper that couples helicase activation with physical stalling to enforce tumour-suppressive alternative splicing programmes.

## Main

Pre-mRNA splicing to remove introns from nascent transcripts is a fundamental step in eukaryotic gene expression that is regulated to control production of alternative spliced isoforms^1–3^. Intron excision and exon ligation are catalyzed by the spliceosome, a multi-megadalton RNA–protein complex that assembles onto each intron from five small nuclear ribonucleoproteins (snRNPs: U1, U2, U4, U5, and U6)^4–8^. Spliceosome assembly is initiated by the U1 snRNP recognizing the 5′ splice site (5′SS) and the U2 auxiliary factor U2AF1/2 binding the 3′ splice site. This early complex is then joined by the U2 snRNP which engages the branch site (BS) to form the prespliceosomal A complex. Subsequent recruitment of the U4/U6.U5 tri-snRNP and the addition and subtraction of multiple other components successively generate the precursor spliceosome (pre-B complex), the precatalytic spliceosome (B complex), the activated spliceosome (B^act^ complex), and the B* complex where the first transesterification reaction is catalyzed. This results in cleavage at the 5′ splice site with ligation of the 2′ hydroxyl of the branchpoint A residue to the phosphate at the 5′ end of the intron. The second catalytic step then results in cleavage at the 3′ splice site, with exon ligation and release of the intron in a lariat structure containing the branched nucleotide. During these many steps of spliceosome assembly, specific ATP-dependent DExD/H-box helicases serve to remodel internal RNA interactions and displace proteins, often with dramatic conformational changes^9–12^. The engagement of each helicase is tightly controlled by trans-acting factors. These factors recruit or stimulate the helicases at the right moment to drive a particular assembly step and ensure fidelity of splice site choices.

Hundreds of additional RNA binding proteins in the human genome modulate spliceosome assembly to determine alternative splice site choice and protein isoform expression^13–15^. These regulatory factors control important splicing programmes in many cellular processes, but their mechanisms of action are largely unknown. One such factor is RNA Binding Motif Protein 5 (RBM5), an alternative splicing regulator that controls apoptosis and cell-cycle dependent gene expression and is frequently disrupted in lung and other cancers^16–23^. In vitro studies found that RBM5 does not affect initial splice site recognition but was present in spliceosomal A complexes where it may prevent further maturation on specific introns^24–26^. We recently identified RBM5 as a component of U2/branchsite complexes in vivo and found that this U2 interaction was required for RBM5 splicing regulation^27^. RBM5 contains multiple functional domains, including two RNA Recognition Motifs (RRMs) and two zinc fingers (ZnFs) for RNA binding^28–32^, an OCtamer REpeat (OCRE) domain potentially interacting with the U4/U6•U5 tri-snRNP^26,33^, and a C-terminal Glycine-rich (G-patch) domain that directly binds the DHX15 helicase and stimulates its activity in vitro^34^. These features suggest that RBM5 may recruit or activate U2-associated DHX15 to induce conformational changes within early spliceosomes that alter splicing choices. However, when and how RBM5–DHX15 function together for such surveillance are unknown.

DHX15 (yeast Prp43) has an established role in disassembling the intron lariat spliceosome following the second catalytic step of splicing^35,36^. Emerging evidence also indicates that DHX15 contributes to early splicing decisions^37–42^ and disruption of this function is linked to cancer^43,44^. These studies suggest that DHX15 selectively suppresses the completion of splicing at suboptimal sites. However, the mechanisms of DHX15 targeting to these early complexes and its discrimination between normal and defective splicing events remain unclear.

Spliceosome assembly is initiated and intron excision is often completed during RNA synthesis, while the precursor RNA is still attached to the polymerase on the DNA template within chromatin^45–47^. The transience of spliceosomes and their heterogeneous and dynamic assembly pathway make studies of their properties in vivo very challenging. Most high-resolution mammalian spliceosomal structures have been assembled in nuclear extract onto simple synthetic RNAs^7,8,48^. The relationship of these structures to spliceosomes assembled onto the long heterogeneous introns of nascent RNA is not yet clear. Moreover, the actions of many splicing regulatory factors that strongly influence alternative splicing decisions in vivo are difficult to reconstruct using in vitro splicing approaches. This limits our ability to integrate the known action of these factors in vivo with our increasingly detailed understanding of spliceosome assembly in vitro.

Here, we isolate spliceosomal complexes bound to nascent RNA from cellular fractions of chromatin-associated nuclear material and determine the cryogenic electron microscopy (cryo-EM) structure of a branchpoint-bound spliceosome at 3.3 Å resolution. This precatalytic complex, with closed SF3B1 engaged with the bulged branch site helix and polypyrimidine tract (PPT), reveals the tumour-suppressor–RBM5 bound within the spliceosome architecture. RBM5 activates DHX15 via its G-patch and anchors the helicase downstream of the branchpoint through extensive interactions with the SF3B1 HEAT-repeats. This RBM5–SF3B1–DHX15 tether is further stabilized by U2SURP/SR140. Our findings reveal RBM5 as a regulatory factor acting on the U2 snRNP to block spliceosome assembly at an early stage. These blocked complexes can be disassembled by the activated DHX15 to control splicing at regulated sites. The structures show how loss of function mutations in a tumour suppressor can result in defective splicing regulation in the early spliceosome.

## Results

### Isolation and cryo-EM of a chromatin-derived spliceosome assembly

To capture RBM5-bound spliceosomes engaged with nascent RNA, we isolated complexes from chromatin fractions of human HEK293 cells following a protocol^27^ modified to optimize the yield and buffer conditions for cryo-EM analysis (see methods). Briefly, nuclei were gently lysed with detergent, and the high molecular weight (HMW) material containing chromatin and nascent RNA was pelleted^49^. This pellet was treated with DNase and RNase to release spliceosomal components, and target spliceosomes containing RBM5-FLAG were affinity-purified on anti-FLAG beads (Fig. 1a). The resulting sample contained a uniform set of U2 snRNP proteins (SF3A and SF3B proteins), characteristic of the 17S U2 snRNP^50,51^ but lacking its 3′ domain components: U2A′, U2B″, and Sm proteins (Fig. 1b,c). Parallel isolations using an antibody to the U2 protein SF3A2 yielded a similar complex except lacking RBM5 (Fig. 1b,c), consistent with the limited expression of endogenous RBM5 in the HEK293 cells and similarly weak RBM5 coprecipitation observed in K562 cells^27^. The chromatin-derived RBM5 isolate was subjected to single-particle cryo-EM analysis, unveiling a SF3B-containing structure (Fig. 1d) at 3.3 Å overall resolution (Extended Data Figs. 1 and 2b and Extended Data Table 1). De novo model building was facilitated by AlphaFold prediction and further validated through protein characterization, using biochemical and in vivo assays to assess protein interactions (Methods).

**Fig. 1.**
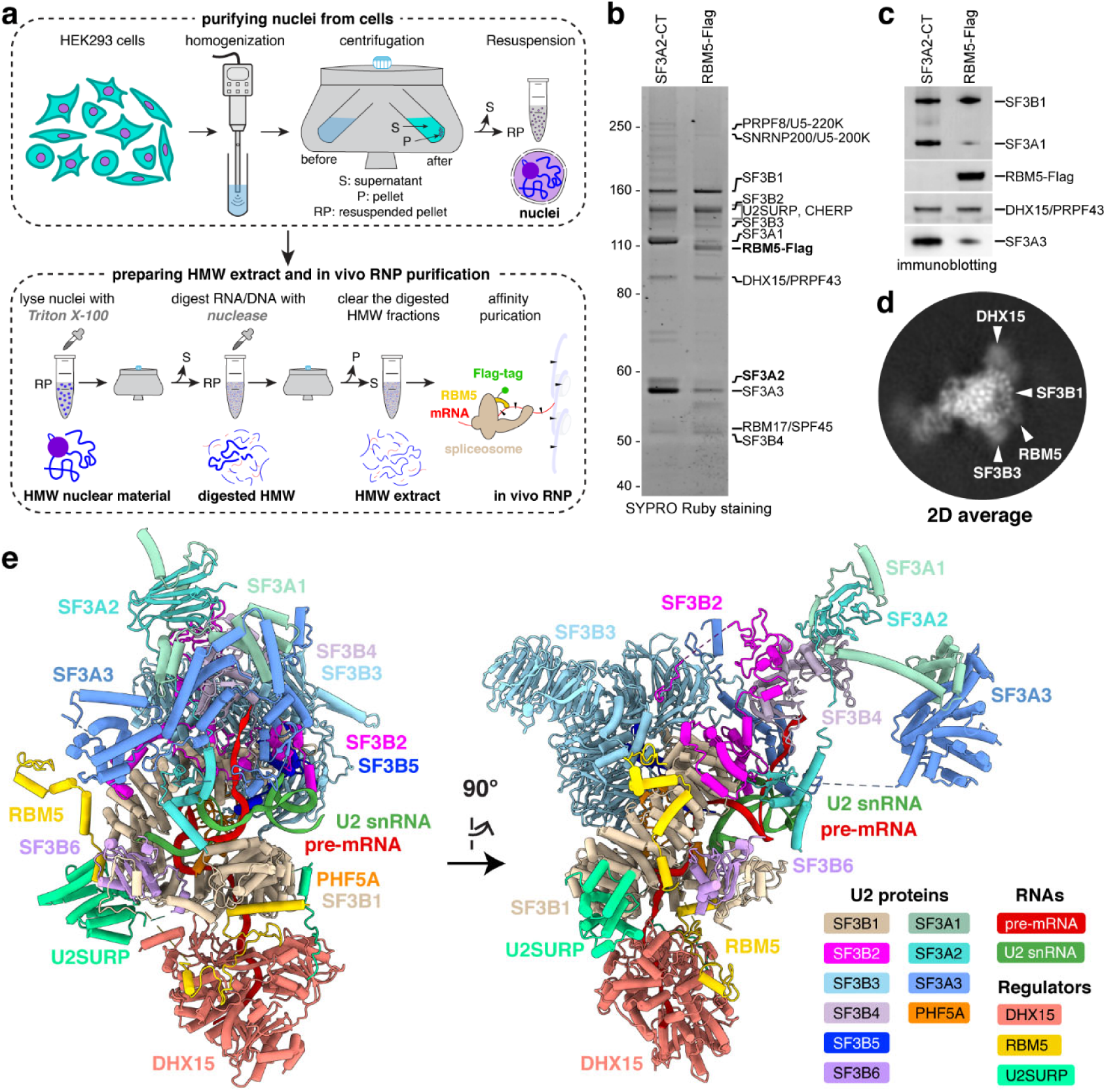
In vivo purification and overall architecture of the chromatin-derived spliceosome. **a,** Workflow for preparing high-molecular-weight (HMW) material from cell nuclei and isolating RBM5-associated ribonucleoprotein (RNP) complexes. Major steps and resulting products are shown at the top and bottom, respectively. **b,c,** In vivo RNP isolation from parental (left) or 293Flp-In cells expressing RBM5-Flag (right) using SF3A2-CT antibody or Flag antibody, respectively. Eluted complexes are separated by SDS-PAGE and visualized by SYPRO Ruby staining (**b**) and immunoblot (**c**). **d,** Representative two-dimensional (2D) class average of the RBM5 spliceosome with features labelled. **e,** Perpendicular views of the atomic model (cartoon representation) of the spliceosomal RNP.

**Fig. 2.**
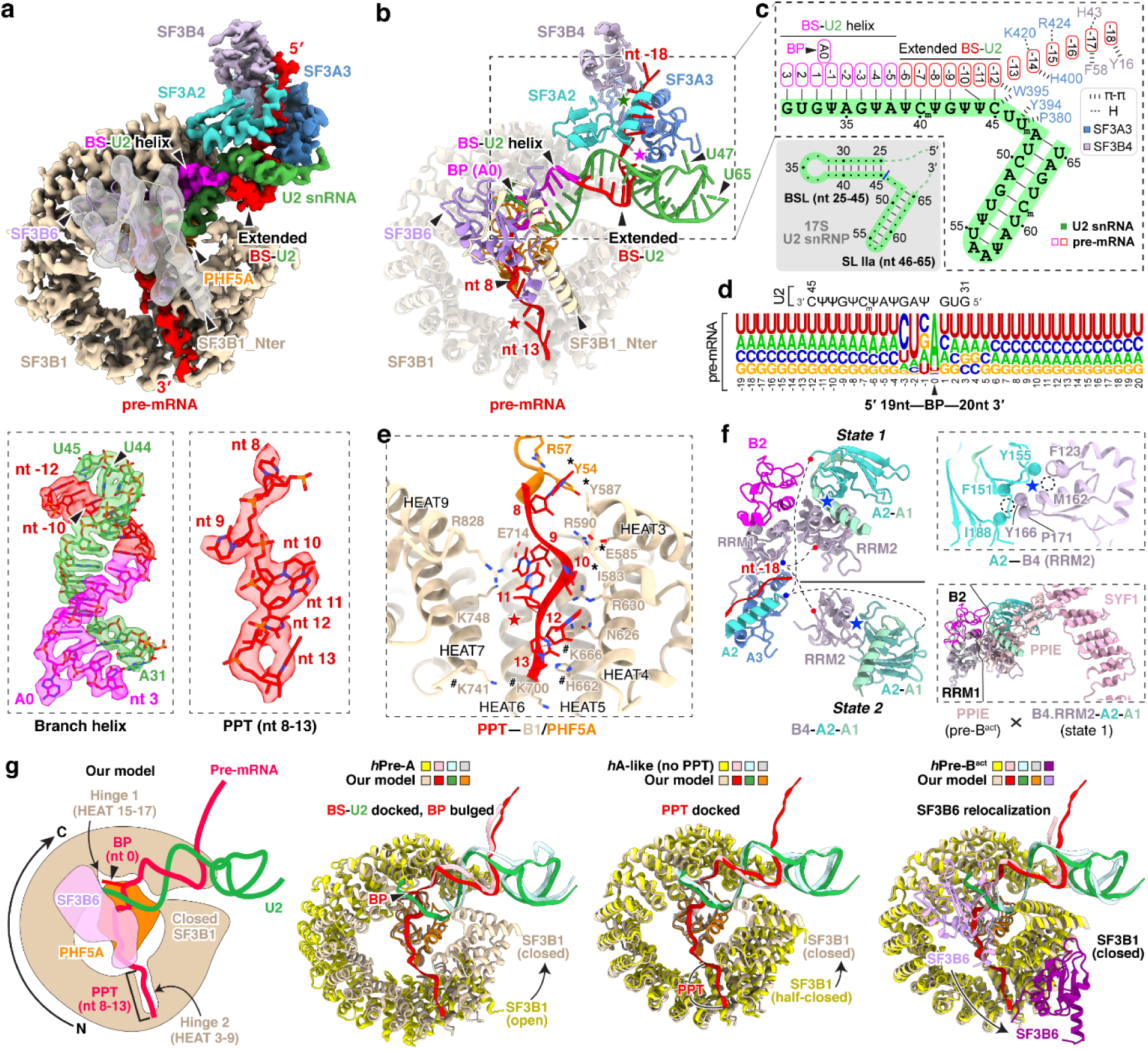
Features of U2/branchpoint (BP) recognition of the in vivo spliceosome. **a,b,** Cryo-EM density maps (**a**) and atomic model (**b**) of the spliceosomal components positioned around the BP. In the top panel of **(a)**: SF3B6 is displayed using a gaussian low-pass filtered transparent map; all others are shown using sharpened densities of the 3.3-Å map. In the bottom panel of **(a)**: the BP adenosine is designated as position 0 of the pre-mRNA, with upstream nucleotides numbered negatively and downstream nucleotides positively. **c,** Secondary structure of the branch site (BS)–U2 snRNA duplex, with nucleotide positions and interacting residues annotated. For comparison, the 2D structure of the human 17S U2 snRNA (PDB: 6Y5Q/7Q4O) is shown at lower left (grey shading), highlighting the branchpoint-interacting stem loop (BSL) and stem loop IIa (SL IIa). **d,** Compilation of the pre-mRNA sequences surrounding the BP bound by the isolated spliceosomal snRNP. The relative letter sizes represent nucleotide frequency at each position. The complementary U2 snRNA sequence is shown above. **e,** Residues engaging the pre-mRNA downstream of BP including the polypyrimidine tract (PPT). Cancer-related mutational hotspot residues are indicated with hash symbols; asterisks denote hydrophobic contacts; labeled residues without symbols contribute to hydrogen bonding. **f,** Two conformational states of the SF3B4–SF3A2–SF3A1 module. Unmodeled regions are shown as dashed lines, with blue and red dots marking the N- and C-termini, respectively. Top right: Identical hydrophobic interactions between SF3B4 RRM2 and SF3A2 in both states. Bottom right: The IBC complex protein PPIE of Pre-B^act-2^ (PDB: 7ABI) clashes with the state 1 conformation. Structures were aligned using the SF3B4 RRM1. **g,** Structural comparison of our U2 snRNP with the reported *human* pre-A (PDB: 7VPX), A-like (PDB: 7Q4O), and pre-B^act-1^ (PDB: 7ABG) spliceosomes. For clarity, only the BS–U2 duplex with surrounding nucleotides, SF3B1, and PHF5A are shown, with SF3B6 included only in the pre-B^act-1^ comparison. A cartoon illustration of our U2 snRNP model is shown in the left panel. Key structural transitions relative to our U2 snRNP are annotated above the right three panels. The middle two panels depict conformational transitions leading toward our structure, whereas the rightmost panel represents a transition away from it.

The overall architecture of the complex included ten U2 snRNP constituents organized into two subcomplexes: SF3B and SF3A (Fig. 1e). The SF3B subcomplex, engaging the branch site, is comprised of seven subunits (SF3B1–SF3B6 and PHF5A) and the U2–branch site helix, collectively forming the 5′ domain of the 17S U2 snRNP. The SF3A subcomplex, which interacts extensively with SF3B, consists of SF3A1, SF3A2, and SF3A3, and contributes to the 3′ domain of the 17S U2 snRNP. Consistent with the SDS–PAGE results and prior proteomic and RNA-sequencing data^27^, the Sm core, U2A′, U2B″, and the U2 snRNA 3′ nucleotides beyond position 65 were not observed, due to RNase-mediated trimming of the solvent-exposed region immediately downstream of U2 stem-loop IIa^51,52^. These findings imply that the isolated U2 exists as part of a larger spliceosomal assembly rather than as an isolated entity. The presence of SF3B proteins indicates that the spliceosome is in a pre-branching state, prior to the B* complex formation^53,54^.

Beyond the canonical U2 proteins, the splicing regulator RBM5—not previously observed in spliceosome structures from nuclear extracts—extensively interacts with the SF3B1 HEAT repeats in this chromatin-derived spliceosome (Fig.1e). Outside of these contacts, regions of RBM5 preceding its second zinc finger (ZnF2), including the two RRMs, ZnF1, and the OCRE domain, were not visible in the structure (Fig.1e), suggesting they may adopt flexible conformations. Of particular note, we identified two additional proteins not previously observed in early spliceosome structures: the DHX15 helicase and U2SURP. Tethered to the RBM5 G-patch domain, DHX15 is situated exterior to the SF3B1 N-terminal HEAT repeats and loaded onto the pre-mRNA segment downstream of the U2–branch site helix. The U2-associated splicing factor U2SURP is seen to engage both the SF3B1 HEAT repeats and DHX15.

### A common U2/branch-site interaction

Unlike previous structures of defined pre-mRNA substrates, our isolated spliceosome was assembled on nascent cellular RNA containing many different introns. This structure presents a common architecture of the intron BS–U2 snRNA duplex (Fig. 2a-c). The structure resolves the bases of the bulged branchpoint adenosine (BP-A, nt 0) and 15 nucleotides of the U2 snRNA that prior to pairing with the BS form the evolutionarily conserved branchpoint-interacting stem–loop in the free 17S U2 snRNP (BSL, nt 31-45) (Fig. 2a, bottom left; Fig. 2c). The presence of this long BS–U2 duplex is striking given the heterogeneity of BS sequences seen in the in vivo sample (Fig. 2d). Upstream of the yUnAy motif enriched at the branchpoint (nt -3 to 1; IUPAC codes: Y = [U, C]; N = [A, U, G, C]), the BS sequences show no base preferences beyond a limited uridine bias (Fig. 2d). Nevertheless, the ensemble of sequences form an extended BS–U2 base paired helical segment similar to spliceosomes assembled onto a single BS sequence (Fig. 2b,c). Each nucleotide from +3 to -9 relative to the BP is paired with its cognate U2 nucleotide (nt 31-42). BS nucleotide -10 pairs with the U2 pseudouridine at nt 44, skipping a pseudouridine at nt 43. (Fig. 2a, bottom left; Fig. 2c).

Downstream of the branchpoint, the pyrimidine preference of the polypyrimidine tract (PPT) before the 3′ splice site is apparent starting at nt +5 (Fig. 2d). The map shows well-resolved density for this segment from nt 8 to 13 (Fig. 2a, bottom right). Apart from nt 8, which contacts PHF5A, the PPT segment is mostly clamped within a cavity formed by SF3B1 HEAT repeats 3-9 (Fig. 2e). These PPT nucleotides were not previously fully defined in human spliceosome structures until the pre-B^act^ complex^55,56^, but in our map the PPT density is clearly resolved within the SF3B1 cavity at this early stage of assembly (Extended Data Fig. 3b, second from left). Positively charged SF3B1 residues K662, K666, K700, and K741, four of the most frequent cancer hotspot mutations^44^, cluster on the inner surface of HEATs 5-7 (Fig. 2e). These residues do not directly contact the 6-nt PPT segment in our structure but are positioned adjacent to its 3′ end (Fig. 2e) where they likely influence PPT placement or binding.

On the opposite side of the BP, the 15-nt BS–U2 duplex forks after C45 of U2 snRNA (Fig. 2c; Extended Data Fig. 4, left). In previous structures, the positions of these residues around the fork were assigned from lower resolution maps. In our map, well-defined density for the duplex terminus and the following U2 nucleotides allowed modeling of U45, U46, and U47 with their interacting SF3A3 residues (Extended Data Fig. 3b, third from left). SF3A3 W395 forms π–π stacks with the terminal bases of the BS–U2 duplex—C45 of U2 and nt -12 of BS—while Y397 and P380 clamp U47 of U2, thus stabilizing the fork in the helix (Extended Data Fig. 4, left). Within the 5′ segment of the pre-mRNA, six nucleotides (nt -18 to -13) beyond the BS–U2 duplex are resolved: nt -14 and -15 contact SF3A3, while nt -16 to -18 interact with SF3B4 RRM1 (Extended Data Fig. 4, right). Notably, SF3A3 H400 stacks with nt -14, and SF3B4 F58 and Y16 π–π stack with nt -17 and -18, respectively (Extended Data Fig. 4, right).

In addition to SF3A3 and SF3B4 RRM1, the region near the 5′ pre-mRNA segment contains SF3A2–SF3A1 and SF3B4 RRM2 that form the SF3A complex region associated with the U2 3′ domain, previously modeled at low resolution by rigid-body fitting with crosslinking data^51,57^. Our map resolves two conformations of the SF3B4 RRM2–SF3A2–SF3A1 complex, termed state 1 and state 2 (Fig. 2f, left; Extended Data Fig. 3c). These share the same SF3B4 RRM2–SF3A2 interface (Fig. 2f, top right) but occupy different positions (Fig. 2f, left), likely reflecting alternative placements of the U2 3′ domain. State 1 adopts a position similar to that in the *human* pre-A complex (Extended Data Fig. 5a), while state 2 resembles the *yeast* pre-A complex (Extended Data Fig. 5b). More precise structural comparison shows our complex differs from both, with relatively smaller deviation from the yeast pre-A (Extended Data Fig. 5c). In a further comparison with the B^act^ stage, alignment of state 1 with pre-B^act-2^ indicates that the IBC protein PPIE would clash with state 1 (Fig. 2f, bottom right), again placing our complex before spliceosome activation.

Other structural features in our complex also help place it along the assembly pathway. In our structure, the branchpoint adenosine is bulged out and stabilized by hinge 1 of a closed SF3B1 together with PHF5A (Fig. 2g, left), an architecture that occurs after the pre-A complex where SF3B1 remains open (Fig. 2g, second from left). SF3B1 stays closed until the B^act^ complex. As seen in our structure, and in other early complexes, SF3B6 is located near PHF5A and the U2/BS duplex, adjacent to C-terminal HEAT repeats of SF3B1. In the pre-B^act^ complex, SF3B6 relocates to the other side of the SF3B1 ring near its N-terminus (Fig. 2g, right). Thus, like the analysis of SF3B4 RRM2–SF3A2–SF3A1, the position of SF3B6 indicates that our complex is at a stage prior to spliceosome activation. Another U2 snRNP conformation seen in an A-like complex shares a similar overall architecture to our structure (Fig. 2g, third from left). This A-like complex was determined with a synthetic RNA lacking a PPT, and—unlike our structure—its SF3B1 adopts a half-closed conformation^58^. Thus, our complex represents an assembly intermediate distinct from earlier structures, and likely formed after the defined pre-A complex but prior to pre-B^act^.

### RBM5–SF3B1 interactions

RBM5 engages extensively with the HEAT-repeats of SF3B1 through two discrete interfaces. At interface I, a helix–loop–helix (HLH) motif of RBM5 binds the convex face of SF3B1 HEAT13–15, predominantly via salt-bridge networks (Fig. 3a, 3b, left). The N-terminal interacting helix (IH1; aa. 686-700) contacts HEAT15 and HEAT14, while the second interacting helix (IH2; aa. 704-711) engages HEAT13 (Fig. 3b, right). Consistent with this, earlier work demonstrated that aa. 544–702 of RBM5, including the HLH motif, are critical for U2 binding^27^. The second RBM5 contact (Interface II) is on the opposite side of the SF3B1 ring from Interface I. Here, the C-terminal helix of RBM5 (C-helix; aa. 799-815), inserts into a groove formed by HEAT2–4 (Fig. 3a, 3b, right), where it establishes a composite interface of both hydrophobic contacts and hydrogen bonds.

**Fig. 3.**
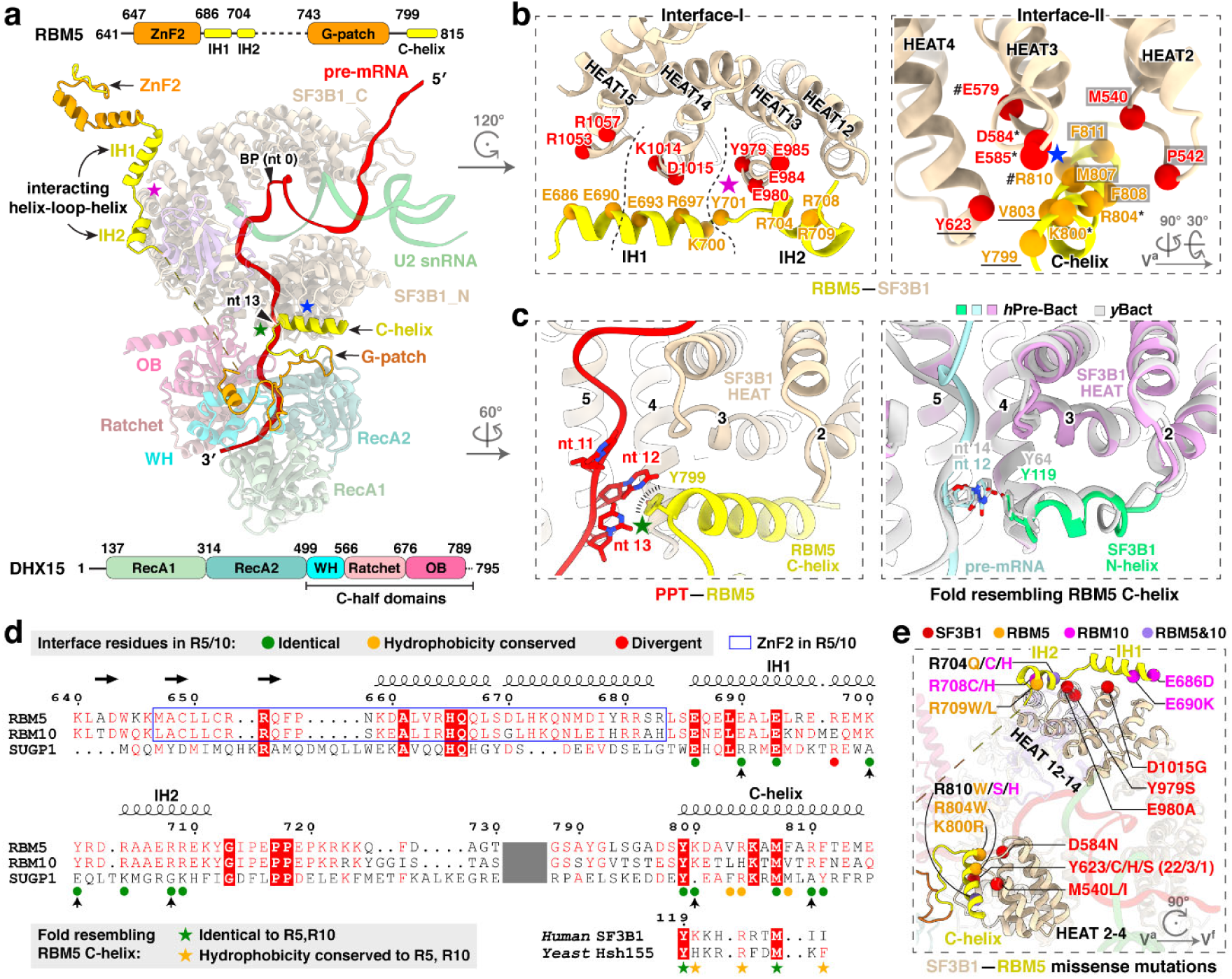
RBM5–SF3B1 interactions. **a,** Domain architecture of RBM5, DHX15, and SF3B1 within the complex. Two RBM5–SF3B1 interfaces are marked with magenta and blue stars, and the RBM5–pre-mRNA contact with a green star. Domain diagrams of RBM5 and DHX15 are shown at the top and bottom, respectively, with residue numbers marking domain boundaries. Unresolved regions of RBM5 are represented by dashed lines (residues 704-743) or omitted (N-terminal residues preceding 641). **b,** Close-up views of RBM5–SF3B1 interaction interfaces, highlighting key binding residues. In Interface-I, three clusters are demarcated by two dashed lines; in Interface-II, four interaction clusters are indicated by hash symbols, asterisks, underlines, and grey boxes, respectively. **c,** The PPT contact with RBM5 C-helix (left) and a matching fold in SF3B1 from B^act^ complexes (right). **d,** Multiple sequence alignment of the G-patch proteins RBM5, RBM10, and SUGP1, showing their C-helices aligned with the SF3B1 N-helix at the bottom. Secondary structure elements from the RBM5 structure are shown above, with β-sheets as horizontal arrows and α-helices as helical coils. Vertical arrows below indicate residues identical in RBM5 and RBM10 but divergent in SUGP1. The omitted sequences are indicated in dark grey. **e,** Single missense mutations from the COSMIC cancer genome database mapped to the SF3B1–RBM5 interfaces.

In addition to its protein contacts, the RBM5 C-helix stacks against a U-base of the PPT through Y799 (Fig. 3c, left), forming a π–π interaction that likely stabilizes the PPT during early spliceosome assembly. In pre-B^act^ structures where the PPT is resolved, an N-terminal helix of SF3B1 occupies the same position as the RBM5 C-helix and forms a nearly identical π–π stack with the PPT (Fig. 3c, right). Both human and yeast SF3B1 preserve residues corresponding to RBM5 C-helix (Fig. 3d, bottom). It appears that this intramolecular SF3B1 contact is mutually exclusive with that of the RBM5 C-helix.

To examine whether other G-patch proteins could make equivalent interactions with SF3B1, we retrieved reported SF3B1–interacting G-patch proteins from the BioGRID database and predicted their structures using AlphaFold (Extended Data Fig. 6a,b). As a positive control, AlphaFold correctly predicted the identified recognition elements of RBM5, including the HLH motif and C-helix that mediate its interaction with SF3B1 (Extended Data Fig. 6b, top left). RBM10 was also predicted to harbor both elements, suggesting that RBM5 and RBM10 engage SF3B1 through shared interactions, consistent with their observed mutually exclusive binding^27^. In contrast, only the C-helix–like element was detected in the predicted structures of SUGP1, CHERP and TFIP11, and no RBM5-like structures were identified in other G-patch proteins (Extended Data Fig. 6b). We examined the SF3B1 contacts with the C-helices of RBM10, SUGP1 and CHERP in AlphaFold, finding that all three helices docked into the same interaction surface of SF3B1. Notably, the C-helix of RBM10 adopted an orientation similar to that of RBM5, whereas the C-helices of SUGP1 and CHERP were oriented in the opposite direction (Extended Data Fig. 6c,d). Consistently, an AlphaFold model of the SF3B1–SUGP1 complex predicts a C-helix orientation similar to that observed here^41^. Consistent with these predictions, the SF3B1-contacting residues were conserved in RBM5 and RBM10 but divergent in SUGP1 and CHERP (Fig. 3d). Within these interfaces, residues of SF3B1, RBM5, or RBM10 have been reported as mutated in cancer genomes of the COSMIC database (Fig. 3e). Although their role in cancer is not yet studied, these contacts of RBM5 with the U2 snRNP presumably contribute to its action as a tumor suppressor.

### DHX15 binds the pre-mRNA exiting SF3B1

A distinctive feature of the RBM5/U2 structure is the presence of the DHX15 helicase, which was previously identified as a component of early spliceosomes^38–40,42^ but has not been resolved in their structures. In our complex, the helicase is bound to the RBM5 G-patch and aligns with HEAT4–9 of SF3B1 (Figs. 3a and 4a). Structural comparisons of our DHX15–G-patch–RNA assembly to other DHX15 structures (G-patch–bound^59^ or RNA-ATP–bound^60^) show the helicase in a more open conformation distinct from these known states (Fig. 4b). Moreover, comparison with the SF3B1-associated helicases DDX42 and DDX46^61^ that act before A-complex formation shows that DHX15 occupies a distinct position on SF3B1 contacting partially overlapping but mutually exclusive surfaces, consistent with DHX15 acting at a later stage of spliceosome assembly (Fig. 4c,d).

**Fig. 4.**
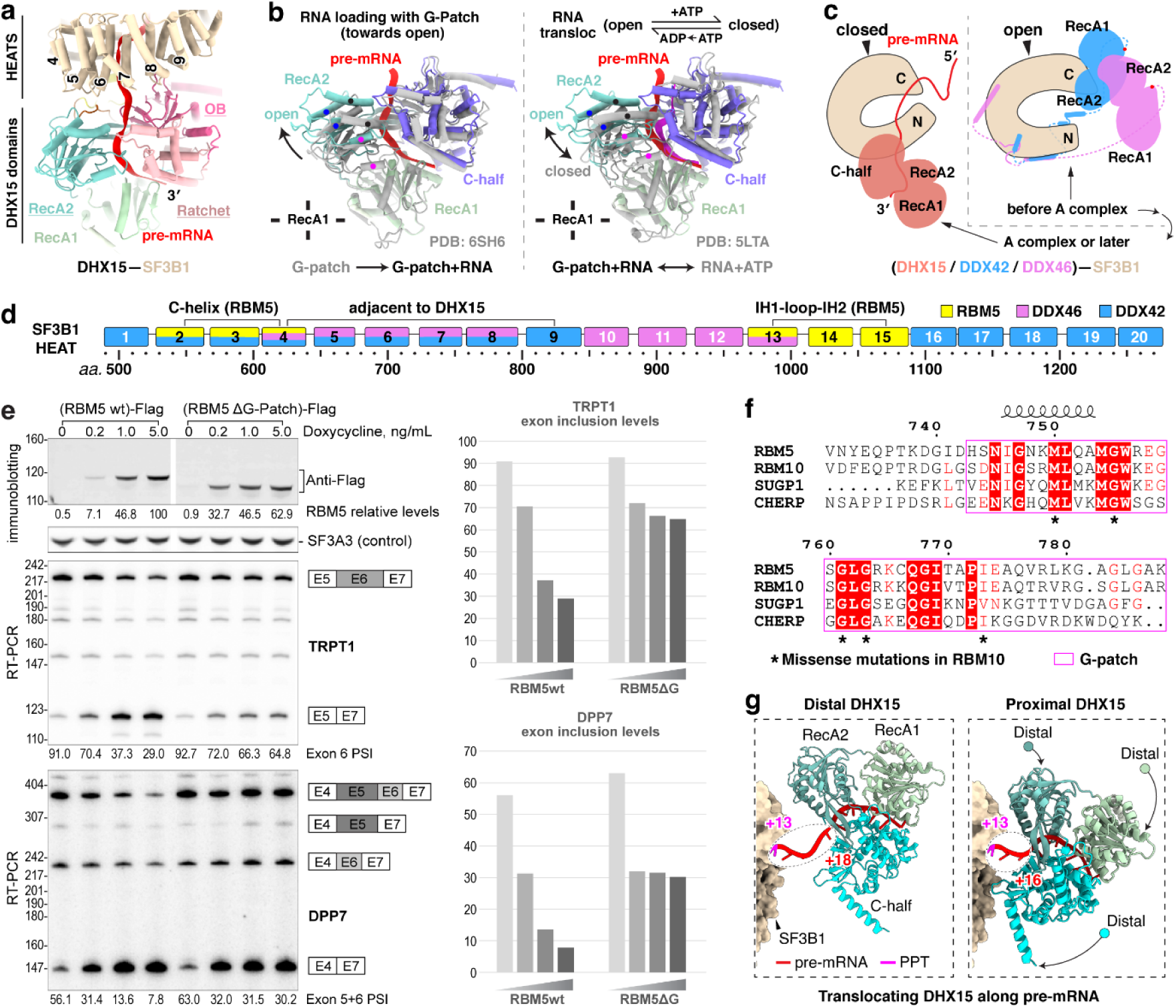
DHX15 is anchored by RBM5 to the 3’ pre-mRNA as it exits SF3B1. **a,** DHX15 positioning near HEAT repeats 4-9 of SF3B1. **b,** Comparison of DHX15 structures in distinct conformational states. Structures are superimposed via the RecA1 domain. Three pairs of dots on α-helices mark their corresponding positions in the two structures to highlight the conformational shift. Left: our DHX15 bound to both G-patch and RNA superimposed with the DHX15 structure when bound only to a G-patch; Right: our DHX15 superimposed with DHX15 bound to both RNA and ATP. **c,** Different SF3B1 binding sites for the human U2 snRNP helicases DHX15, DDX42, and DDX46 (yeast Prp5 homolog). The SF3B1 N-terminal and C-terminal regions are labeled as N and C, respectively. Unresolved helicase regions are shown as dashed lines, with red dots marking their C-terminal ends on the helicase C-terminal half. **d,** Diagram of SF3B1 interactions with RBM5 and RNA helicase interactions along the SF3B1 HEAT repeats. **e,** The RBM5 G-patch domain promotes exon skipping in TRPT1 and DPP7 pre-mRNA. <u>Top left</u>: Expression levels of Flag-tagged RBM5 and RBM5 ΔG-patch, quantified by immunostaining, with SF3A3 as a normalization control. <u>Middle and bottom left</u>: RT-PCR analyses of endogenous TRPT1 exon 6 (middle) and DPP7 exons 5+6 (bottom) splicing, using radiolabeled flanking primers, followed by denaturing PAGE and phosphorimaging. Quantified exon inclusion levels are displayed below each RT-PCR result. <u>Right</u>: Graph of quantified exon inclusion levels. **f,** Multiple sequence alignment of the G-patch proteins RBM5, RBM10, SUGP1 and CHERP. Secondary structure elements from the RBM5 structure are shown above with α-helices as helical coils. Asterisks below indicate RBM10 residues subject to single missense mutations from the COSMIC cancer genome database. **g,** Structures showing two translocation states of DHX15 along pre-mRNA, with nucleotide numbers downstream from the branch point (BP) adenosine.

To specifically assess the significance of the DHX15–SF3B1 tether in RBM5 function, we deleted the G-patch from RBM5 and expressed this protein in HEK293 cells at different levels by titrating its induction with Doxycycline. This delta G-patch RBM5 was previously found to maintain binding to the U2 snRNP^27^. We monitored the splicing of multiple known RBM5 target exons in the presence of either wildtype or G-patch-deleted protein. For exons in the TRPT1 and DPP7 transcripts, the G-patch deletion mutant showed reduced activity compared to the wildtype RBM5. Interestingly, the G-patch mutant protein induced partial exon inhibition at low expression but this was the limit of its activity. In contrast to the wildtype protein, no additional reduction in target exon splicing was observed with higher concentrations of mutant RBM5 (Fig. 4e). These observations suggest that RBM5 may induce some level of exon repression even without its G-patch. On other RBM5 downregulated exons including NUMB exon 9, the activity of the G-patch mutant was comparable to that of the wildtype RBM5 (Extended Data Fig. 7). Together, these results imply there are two distinct but not mutually exclusive pathways of RBM5-mediated exon inhibition, one mediated by the G-patch and one not.

When tethered to SF3B1 via the RBM5 G-patch, DHX15 is bound to the 3′ pre-mRNA sequence as it exits the U2 snRNP—where DExH-box translocation is predicted to disrupt upstream RNA-protein interactions. Through further particle classifications, we identified two distinct DHX15 states: a distal state with the protein located toward the 3′ end in the pre-mRNA and farther from SF3B1, and a proximal state where DHX15 engages a more upstream RNA segment and is positioned closer to SF3B1 (Fig. 4g). Together, these states reveal a DHX15 translocating along the pre-mRNA. Notably, DHX15 does not form discernible protein contacts with SF3B1, even in the proximal state, indicating that its placement is likely determined by the RBM5–SF3B1 association rather than direct interaction with SF3B1.

### U2SURP tethers RBM5, SF3B1 and DHX15

In addition to RBM5 and DHX15, three conserved segments of U2SURP are resolved in our structure, engaged with the SF3B1 HEAT repeats and DHX15 (Fig. 5a; Extended Data Fig. 8). First, U2SURP residues 534–679 form a CID-like domain that shares 21% sequence identity with the canonical CID (PDB 4NAC) and aligns with the canonical domain with an RMSD of 1.9 Å over 131 residues (Fig. 5b). In our structure, this domain contacts SF3B1 HEAT10–13 and RBM5 IH2 (Fig. 5c). Second, an N-terminal segment of U2SURP forms a helix that associates with SF3B1 HEAT4–5 primarily through hydrophobic interactions (Fig. 5d, left). Finally, residues 136–141 of U2SURP form main-chain contacts with the RecA2 β-sheet of DHX15 (Fig. 5d, right). These latter two elements (residues 115–142) constitute an SF3B1–DHX15 bridging module (SDBM). Together, these interactions allow U2SURP to serve as a scaffold that facilitates the association of RBM5, DHX15 and SF3B1 in this early spliceosome. The COSMIC catalog of cancer somatic mutations reports 26 single-missense variants in the CID (Fig. 5c). Although their pathogenicity has not been studied, these include three residues in the CID SF3B1/RBM5 interfaces.

**Fig. 5.**
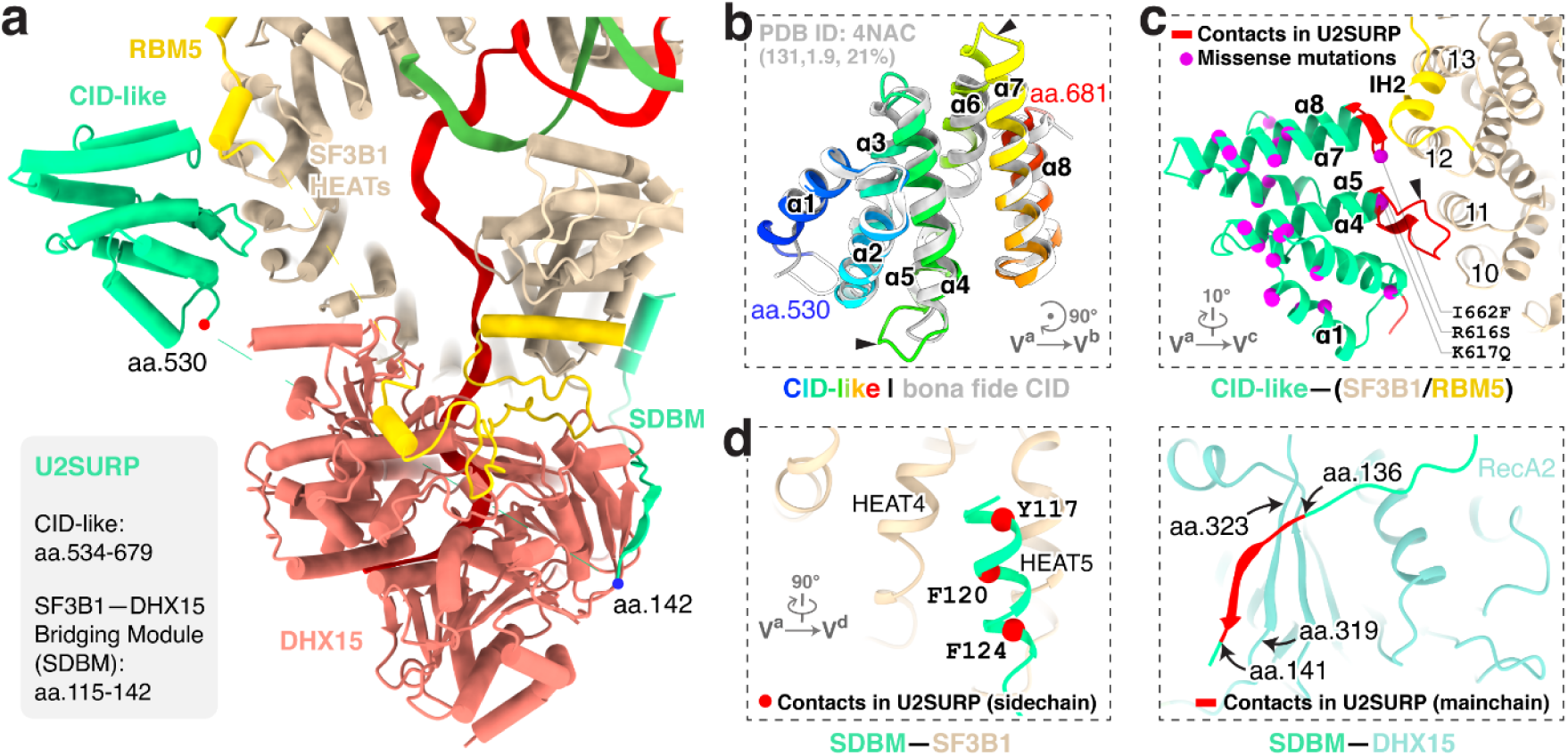
U2SURP promotes RBM5–SF3B1–DHX15 tethering. **a,** Overall view of U2SURP interacting with SF3B1, RBM5 and DHX15. **b,** Domain architecture of the U2SURP CID-like domain and its superposition with a bona fide CID (PDB ID: 4NAC). The three numbers in brackets indicate, from left to right, the number of aligned residues, the Cα r.m.s.d., and the percent sequence identity. **c,** Interaction of the U2SURP CID-like domain with SF3B1 and RBM5. Interacting regions in U2SURP are colored in red. COSMIC single missense alterations mapped on our CID-like structure are shown as magenta spheres. **d,** U2SURP SDBM contacts on SF3B1 (left) and DHX15 (right). U2SURP interactions via side chains (left) and the main chain (right) are shown in red with SF3B1 and DHX15, respectively.

### Stage of the RBM5-stalled spliceosome

Our structure reveals a defined PPT within a closed SF3B1 and with the RBM5 C-helix wedged at HEAT repeats 2–4 (Fig. 6c; Extended Data Fig. 9, middle). Consistent with this, the PPT-absent A-like complex^58^ and PPT-detached B^AQR^ complex^62^ adopt half-closed and loosened SF3B1 conformations, respectively (Fig. 6c, left; Extended Data Fig. 9, left and right), both lacking the RBM5-binding grove (Fig. 6c, right). Notably, in the B^AQR^ complex, the PPT is engaged by Prp2 (Extended Data Fig. 9, right), which adopts a DExH-helicase configuration clearly distinct from the SF3B1-associated DHX15 in our structure.

**Fig. 6.**
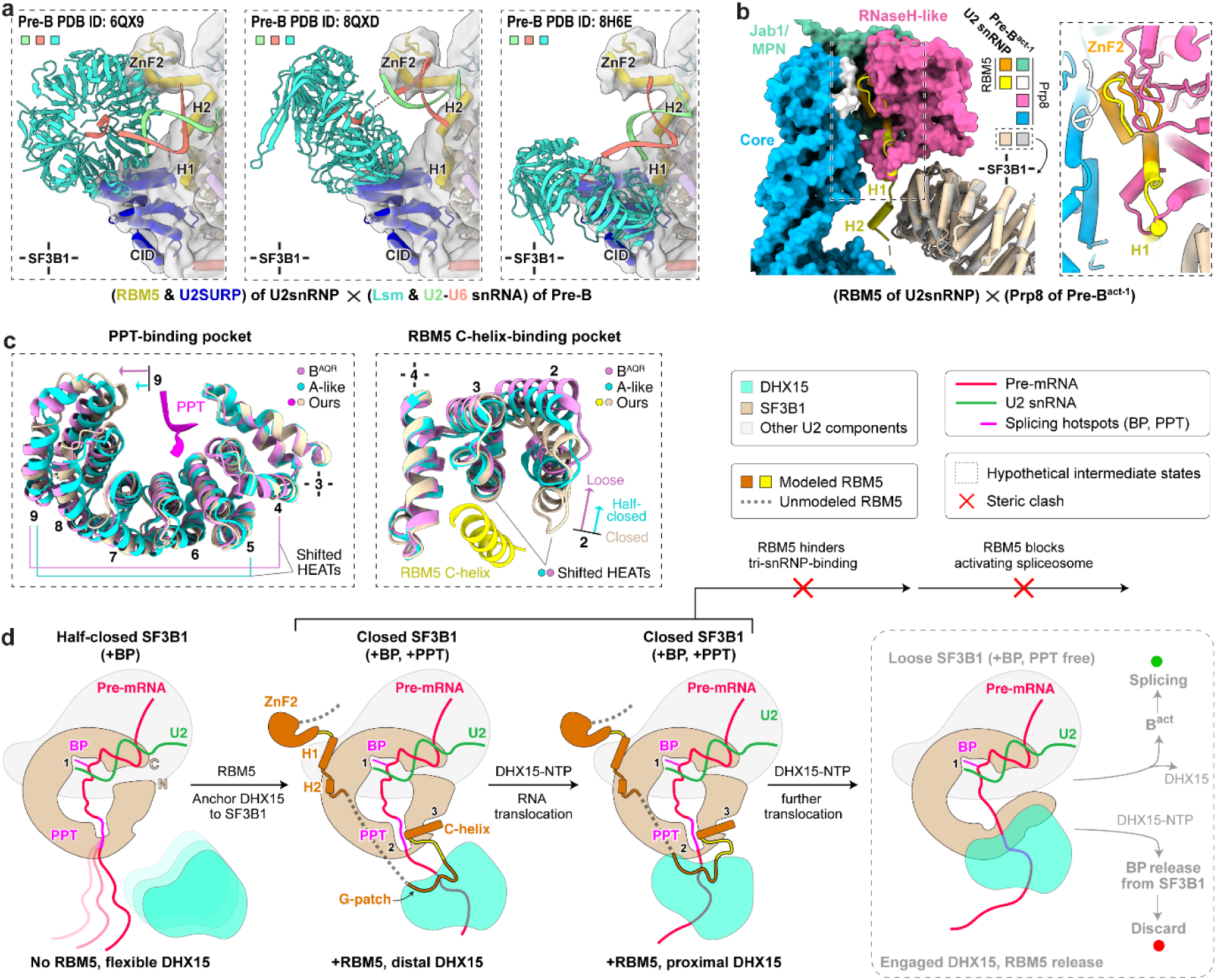
Effects of RBM5-DHX15 binding on spliceosome assembly. **a,** Steric clashes between RBM5/U2SURP and components of three pre-B complex structures. The Lsm ring and the U2-U6 snRNA duplex in pre-B are shown as ribbons, while RBM5 and U2SURP are shown as cylinder cartoon within the density indicated in transparent grey. **b,** Steric clashes between RBM5 and Prp8 of the pre-B^act-1^ complex. Prp8 in pre-B^act-1^ is shown as surface (left) and close-up cartoon (right), while RBM5 is depicted as yellow or orange cartoon on both sides. Structures are superimposed using SF3B1. **c,** Structural rearrangements of the C-helix and PPT-binding pockets in SF3B1 from the closed conformation (our model) to the half-closed (A-like, PDB: 7Q4O) and loose states (B^AQR^, PDB:7QTT). Arrows indicate conformational shifts relative to our model. HEAT repeats used for structural superimposition are marked with cross. **d,** Schematic model for branch site (BP) rejection mediated by RBM5–DHX15.

To place our complex in the sequence of spliceosome assembly, we compared its structure with those of other assembly intermediates. The ZnF2 and HLH motifs of RBM5, together with the CID-like domain of U2SURP, show steric clashes with the Lsm proteins and the U2–U6 snRNA duplex in human pre-B complexes (Fig. 6a), placing our complex prior to the pre-B stage. A severe clash between the RBM5 HLH motif and Prp8 in the pre-B^act-1^ complex is also consistent with this assignment (Fig. 6b). Together with the SF3B1 conformation indicating a state after the pre-A or A-like complexes (Fig. 2g), our structure is best assigned as a prespliceosomal A complex. Unlike the canonical A complex, the presence of RBM5 and the predicted activity of DHX15 in its observed position are predicted first to block further assembly and then to disrupt the U2 interaction thereby precluding productive splicing.

## Discussion

We determined the structure of an in vivo-assembled U2/branchpoint complex bound to the regulatory protein RBM5. Assembled on nascent RNA from the chromatin fraction of cells, the sample contains an ensemble of complexes with many different intron sequences, yet these complexes are remarkably homogeneous in structure and the pre-mRNA backbone can be traced accurately from its entrance into the U2 snRNP to its exit from SF3B1. The observed interactions of RBM5 and its associated DHX15 provide two mechanisms for how this regulator negatively affects spliceosome assembly. The architecture closely resembles the prespliceosomal A complex, but the positions of the RBM5 ZnF2 and HLH domains, and the U2SURP CID domain—all in contact with the SF3B1 ring—block docking of the incoming U4/U6.U5 tri-snRNP and preclude progression to a pre-B complex. Moreover, tethering of DHX15 to SF3B1 by RBM5 and U2SURP, and its loading onto the 3′ pre-mRNA segment as it exits the U2 complex, enable disruption of the complex by the helicase.

One mechanistic model for splicing repression by RBM5 is illustrated in Figure 6d. Prior to the A complex, RBM5 may not be bound and the architecture may resemble the reported A-like state^58^, with SF3B1 half-closed and lacking a defined PPT pocket. Biochemical studies indicate DHX15 is associated, but its lack of visibility suggests flexible tethering (Fig. 6d, leftmost panel). As the complex transitions to the closed state, RBM5 associates with SF3B1 HEAT repeats via the HLH motif and C-helix, and the C-helix stabilizes the PPT within closed SF3B1. DHX15, bound to the RBM5 G-patch, initially engages the 3′ pre-mRNA in a distal position (Fig. 6d, second from left). Upon NTP hydrolysis, DHX15 translocates and anchors at the SF3B1 exit (Fig. 6d, third from left). RBM5 binding interferes with tri-snRNP engagement—producing steric clashes with Lsm proteins, the U2–U6 duplex, and core B^act^ components—thereby stalling progression. After arresting assembly, RBM5 enables continued DHX15 translocation to ultimately disrupt the PPT–SF3B1 contacts and/or detach U2 from the branch site.

Reverse trajectories are also plausible: RBM5 binding could displace tri-snRNP components from a pre-B complex, or DHX15 movement could dislodge the PPT, opening SF3B1 and destabilizing RBM5 association, allowing normal splicing to resume. In this scenario DHX15 would counteract RBM5-mediated repression.

Although we favor DHX15 acting in negative regulation with RBM5, there is evidence that DHX15 may oppose RBM5 repression. Both IP-seq and CLIP analyses show RBM5 binding to unregulated branch sites^27^. This promiscuous binding may be tolerated because DHX15 can disrupt the RBM5–SF3B1 interaction and permit forward assembly. The isolated complex may thus represent a reversible checkpoint that can either proceed to productive splicing or be discarded. Alternatively, it may already be terminally stalled and poised for disassembly. Notably, the G-patch is required for repression of only a subset of RBM5-dependent exons (Fig. 4e, Extended Data Fig. 7), leaving open whether residual helicase activity or simple steric blocking accounts for the remaining repression.

DHX15 was recently implicated in early spliceosome assembly via SUGP1^39–41,63^ and GPATCH8^42^, but those interactions and target complexes remain unresolved. We find that the RBM5 G-patch activates DHX15 and its flanking α-helices orient the helicase at the pre-mRNA exit of the prespliceosomal A complex (Fig. 3a,b). RBM10 has equivalent flanking helices, while SUGP1 and CHERP have C-terminal helices predicted to dock in opposite orientation onto the same SF3B1 region (Extended Data Fig. 6b,c). Interestingly, early studies showed that CHERP and RBM17 form a trimeric complex with U2SURP^64,65^; all three of these proteins are present in our isolated complex but only U2SURP segments are visible (Fig. 1b,e), suggesting they may orient DHX15 in other assembly states. Thus, this SF3B1 segment serves as a common docking platform for multiple G-patch proteins and DHX15.

Finally, our findings illuminate splicing misregulation in cancer. RBM5 loss alters the splicing of target exons affecting apoptosis and other oncogenic pathways^21,23,26,30,66,67^. Well known cancer-associated mutations in SF3B1 also impair DHX15 recruitment by other G-patch proteins^42,63^. We find that numerous missense mutations in the COSMIC catalog map precisely to the RBM5–U2SURP–DHX15–SF3B1 interfaces defined in our structures (Figs. 3e, 4f, and 5c). The disruption of these interactions will alter spliceosome assembly and remodeling with consequences for oncogenic splicing programmes.

## Methods

### Isolation of the chromatin-derived spliceosomal complex

The complexes were purified from HMW extracts as previously described^27^ with the following modifications. 293 Flp-In cells carrying a RBM5–FLAG gene were seeded into thirty-six15-cm plates in the presence of reduced concentration of doxycycline (5 ng/mL) and grown to confluency for three days. Cells were harvested in PBS buffer and homogenized in low salt, high sucrose buffer as above. Cell nuclei were pelleted through a high sucrose cushion. The nuclear pellet was lysed in ten volumes of isotonic lysis buffer containing Triton X-100, the high-molecular weight material was pelleted and the soluble nucleoplasm discarded. Enzymatic extraction was performed at 23–25 °C as above but completed within 45 min. The RBM5/branch site complex was affinity-purified directly from a cleared extract instead of from glycerol gradient fractions to increase yield. Complexes were immunoprecipitated overnight at 4 °C using 20 μl packed M2 Flag–agarose (Millipore-Sigma, A2220), washed four times with buffer containing 20 mM HEPES–KOH pH 7.9, 150 mM NaCl, 1.5 mM MgCl₂, and 0.05% Triton X-100, followed by a final wash in detergent-free buffer. The complexes were eluted on a Thermomixer (15 s at 900 rpm followed by 4 min 45 s rest per cycle) for 2 h at 5 °C in 80 μl of buffer containing 20 mM HEPES–KOH pH 7.9, 150 mM NaCl, 1.5 mM MgCl₂, 10 mM DTT, 4.5% glycerol, and 150 ng/μl 3×FLAG peptide (Millipore-Sigma, F4799).

### Protein analysis

Immunopurified proteins were separated by SDS-PAGE and detected by immunoblotting with primary antibodies^27^ and fluorescently-labeled secondary antibodies (GE Healthcare Bio-Sciences), or stained with SYPRO Ruby (Thermo Fisher Scientific). Images were visualized on an Amersham Typhoon and quantified as above.

### Pre-mRNA sequence analysis

RBM5-Flag RNPseq reads^27^ from intronic 3’ splice site clusters of HEK293 cells were aligned at their predicted branchpoint nucleotides and trimmed 19nt upstream and 20nt downstream. Trimmed reads shorter than 40nt were discarded. Nucleotide frequencies in a random subset of 15,000 trimmed reads were plotted with WebLOGO^68^.

### Electron microscopy (EM) of both stained and frozen-hydrated samples

Negative stain EM was performed by applying 2.5 µl of the chromatin-derived spliceosomal complex to a glow-discharged carbon-coated grid. After a 30 s adsorption, grids were stained with 2% (w/v) uranyl acetate. Micrographs were acquired on an FEI Tecnai F20 electron microscope operated at 200 kV, using a TIETZ F415MP 16-megapixel CCD camera at a nominal magnification of 70,000×.

For cryo-EM sample optimization, an aliquot of 3 μL of sample was applied onto a glow-discharged lacey grid coated with thin continuous carbon (400 mesh, Ted Pella) for 60 s. The grid was blotted with Grade 595 filter paper (Ted Pella) and flash-frozen in liquid ethane with an FEI Mark IV Vitrobot. The same FEI TF20 instrument and imaging condition as negative staining evaluation were used to screen cryo-EM grids. The grids with optimal particle distribution and ice thickness were obtained by varying the gas source (air using PELCO easiGlowTM, target vacuum of 0.37 mbar, target current of 15 mA; or H2/O2 using Gatan Model 950 advanced plasma system, target vacuum of 70 mTorr, target power of 50 W) and time for glow discharge, the volume of applied samples, chamber temperature and humidity, blotting time and force, as well as drain time after blotting. Our best grids were obtained with 30 s glow discharge using air and with the Vitrobot sample chamber set at 8 °C temperature, 100% humidity, 10 s blotting time, 10 blotting force, and 0 s drain time.

For cryo-EM sample optimization, 3 µl of sample was applied to glow-discharged lacey carbon grids with a thin continuous carbon support (400 mesh; Ted Pella) and incubated for 60 s. Grids were blotted with Grade 595 filter paper (Ted Pella) and plunge-frozen in liquid ethane using an FEI Mark IV Vitrobot. Screening was performed on an FEI Tecnai F20 operated under the same imaging conditions used for negative-stain evaluation. Optimization was achieved by varying the glow-discharge gas (air using a PELCO easiGlow™ at 0.37 mbar and 15 mA; or H₂/O₂ using a Gatan Model 950 at 70 mTorr and 50 W), discharge duration, chamber temperature and humidity, blotting time and force, and post-blot drain time. The best ice and particle distributions were obtained using a 30 s air glow discharge and Vitrobot settings of 8 °C chamber temperature, 100% humidity, 10 s blotting time, blot force 8, and 0 s drain time.

Optimized cryo-EM grids were loaded into an FEI Titan Krios microscope equipped with a Gatan Imaging Filter (GIF) Quantum LS and a post-GIF K3 Summit direct electron detector. Data were collected at 300 kV with a GIF slit width of 20 eV. Movies were acquired in super-resolution electron-counting mode using SerialEM^69^ at a calibrated pixel size of 0.55 Å (K3) with a total exposure of ∼50 e⁻ Å⁻² per movie. The movies were acquired by image shift combined with multi-shot collection in SerialEM. Data acquisition parameters are provided in Extended Data Table 1.

### Structure determination

Frames from each movie were aligned for drift correction using MotionCor2^70^. The first frame was discarded to minimize the effects of initial drift and charging. For each movie, two corrected micrographs were generated—one with dose weighting and one without. The aligned micrographs had a calibrated specimen-scale pixel size of 1.1 Å. Non–dose-weighted micrographs were used solely for CTF estimation, whereas dose-weighted micrographs were used for all subsequent image-processing steps.

The defocus value of each averaged micrograph was determined by CTFFIND4^71^ to be ranging from -1.5 to -3 μm. Initially, particles were automatically picked from 20,795 curated averages using both Gautomatch (https://www2.mrc-lmb.cam.ac.uk/research/locally-developed-software/zhang-software/) and cryoSPARC’s blob picker^72^. Gautomatch particles were boxed at 384 × 384 pixels and binned to 192 × 192 pixels (2.2 Å per pixel) before processing in RELION^73,74^, whereas blob-picked particles were processed in cryoSPARC. Multiple rounds of reference-free 2D classification were performed to remove particles in poorly resolved or non-particle classes (junk, dissociated particles, and other contaminants). High-quality class averages were used for cryoSPARC template picking, and the corresponding good particles were also used to train Topaz models, followed by rounds of 2D-based screening. In total, 1,096,950 particles were retained from all these 2D classification steps.

The 1,096,950 particles were subjected to *ab* initio reconstruction in cryoSPARC. The class displaying good model features—intact features as shown in representative 2D classes plus visible secondary structural elements like α-helices—was selected and used as the initial model for a three-class 3D classification in RELION, yielding 513,604 particles in the best class. <u>To obtain a high-resolution map (processing branch 1)</u>, the particles were subjected to three parallel 2D classifications and a duplicate removal, yielding 391,630 unique particles. Three cycles of “ab initio reconstruction and heterogeneous refinement” in cryoSPARC selected 128,413 particles. After unbinning, Non-Uniform Refinement in cryoSPARC produced a 3.40 Å map. Local and global CTF refinements—per-particle defocus and per–exposure-group beam tilt, trefoil and anisotropic magnification—were then applied, followed by a final Non-Uniform Refinement that reached 3.26 Å (gold-standard FSC). <u>To analyze DHX15 (processing branches 2 and 3),</u> the 513,604 particles were subjected to an additional three-class 3D classification in RELION, yielding 308,155 particles in the best class. To minimize the influence of the flexible SF3A region during 3D classification, a soft mask over the SF3A density was applied using the solvent_mask2 option in a 3D classification in RELION. Two classes displaying distinct DHX15 conformations were selected. Particles from each class were unbinned and refined independently by a cycle of Non-Uniform Refinement, CTF refinement and a final Non-Uniform Refinement, producing maps at 3.43 Å and 3.93 Å, respectively. Local classification of the DHX15 region in the 3.43 Å reconstruction further improved its density and revealed secondary-structure features.

Alternatively, <u>to analyze the SF3A region (processing branches 4 and 5),</u> the 1,096,950 particles were subjected to a five-class 3D classification in RELION, from which two classes showing distinct SF3A conformations were selected. For the conformation represented by 187,793 particles (branch 4), a subsequent three-class 3D classification in RELION yielded 138,739 particles in the best class. These particles were unbinned and refined using Non-Uniform Refinement to obtain a 3.80 Å map. Local classification focused on SF3A followed by Non-Uniform Refinement improved the density, and CTF refinement with a final Non-Uniform Refinement further enhanced the resolution to 3.61 Å. Applying a similar procedure to the second SF3A class generated a 3.94 Å reconstruction (branch 5). The full data-processing workflow is summarized in Extended Data Fig. 2a.

### Resolution assessment

All resolutions reported above are based on the “gold-standard” FSC 0.143 criterion^75^. FSC curves were calculated with soft masks, and high-resolution noise substitution was applied to correct for mask-induced convolution artifacts^75^. Density maps were sharpened by application of an automatically determined negative B-factor^76^. Local resolution was estimated in cryoSPARC using the Local Resolution Estimation tool with a local FSC threshold of 0.5. Representative map quality is shown in Extended Data Fig. 2b–f, and data collection and reconstruction statistics are summarized in Extended Data Table 1.

### Model building and refinement

To guide model building of the chromatin-derived RNP, the U2 portions of published structures—from the 17S U2 snRNP to the B^act^ complex—were rigidly docked into our cryo-EM map (Extended Data Fig. 2a, the final map from branch 1) using ChimeraX^77^ to assess their fit. The A-like structure (PDB 7Q4O) provided the best agreement and was selected as the initial model, defining the U2 snRNP configuration as an A, pre-B or B-like state. This model further revealed three peripheral extra densities in our reconstruction (Extended Data Fig. 2a, final maps of branches 2 and 3): one adjoining the 3′ pre-mRNA exit of SF3B1 and positioned near SF3B1’s HEAT repeats 4–9; a second associated with the outer concave surface of SF3B1 HEAT repeats 10–15; and a third located near the 5′ region of the pre-mRNA and the SF3B4 RRM1–SF3B2 heterodimer.

To interpret the remaining extra densities, we examined AlphaFold3^78^-predicted structures of all proteins present in the SDS–PAGE of the purified RNP (Fig. 1b). DHX15 was first rigidly docked into the extra density adjacent to SF3B1 HEAT repeats 4–9 (Extended Data Fig. 2a, final map of branch 2), consistent with its DExH-helicase positioning near 3′ RNA. A G-patch element was then assigned based on the structural arrangement of DHX15–G-patch complexes (PDB 6SH6), with flanking segments contacting SF3B1 HEATs 2–4 and 12–15. AlphaFold3 predictions for the G-patch–containing proteins identified in our gel further supported the assignment of the second extra density to the RBM5 ZnF2 domain and its downstream helix–loop–helix motif, in agreement with previous findings that the region between ZnF2 and the G-patch mediates RBM5–SF3B1 binding^27^. This assignment was reinforced by high-confidence AlphaFold3 predictions of the RBM5 C-helix (aa. 799-815) interacting with SF3B1 HEATs 2–4. The remaining portion of the second density was attributed to the U2SURP CID domain (aa 530–681), again supported by AlphaFold3 predictions and the SDS–PAGE profile. An N-terminal segment of U2SURP (aa 115–142) was also tentatively identified interacting with both SF3B1 and DHX15, guided by the AlphaFold3 modeling and our cryo-EM density. For the third extra density, the 17S U2 snRNP structure (PDB 6Y5Q) provided an initial reference, as it contains a large SF3A-associated density located near—but not matching—the position observed in our reconstruction. The well-resolved density in the locally classified maps (Extended Data Fig. 2a, final maps of branches 4 and 5) allowed confident modeling of the SF3B4 RRM2–SF3A2 heterodimer, which was further supported by rigid docking of the protruding SF3A module.

RNA modeling was guided by the well-resolved cryo-EM density of the BS–U2 duplex and adjacent pre-mRNA on both sides. Branchpoint sequences were assigned according to nucleotide-frequency profiles derived from RNA-seq data (Fig. 2d) and therefore represent a consensus rather than a single endogenous RNA sequence. Downstream of the branchpoint, well-defined density for the PPT segment (nt +8 to +13) enabled modeling of the six-nucleotide pyrimidine tract within the SF3B1 HEAT-repeat cavity using side-chain-supported features. On the 5′ side of the pre-mRNA, improved density at the fork following U2 C45 permitted confident placement of U45–U47 and their interactions with SF3A3.

<u>The central regions of the cryo-EM map</u>, ranging in resolution from 3.5 to 4.5 Å (see Extended Data Fig. 2e), exhibit sufficient detail for manual tracing and *de novo* model building using COOT^79^. Sequence assignment primarily relied on discernible densities corresponding to amino acid residues with bulky side chains, such as Trp, Tyr, Phe, and Arg (Extended Data Fig. 3). Distinctive patterns within sequence segments containing these residues served as benchmarks for validating residue assignments. <u>The peripheral regions of the cryo-EM map</u> did not reach sufficient resolution for de novo atomic modeling. In the final chromatin-derived RNP model, the following components were therefore built using AlphaFold predictions that were rigidly docked into the density: two conformations of the SF3B4 (RRM2, aa. 98–213)–SF3A2 (aa 110–210)–SF3A1 (aa. 160–235, 248–318)–SF3A3 (aa. 1–350) module, SF3B6–SF3B1.N-terminal segment (aa. 378–424), DHX15, RBM5 (aa. 641–700, 743–798), and U2SURP (aa. 115–142, 530–681). We also tentatively traced the main chain of the stretched SF3A2 loop (aa. 93–109) and the SF3A1 linker (aa. 236–247). In addition, the pre-mRNA segment extending beyond DHX15 and SF3B1 was tentatively modeled: nt 4-7 in all models, nt 14–15 in the proximal-DHX15 state and nt 14–17 in the distal-DHX15 state.

Models were refined in real space using PHENIX^80^ with secondary structure and geometry restraints. Refinement statistics are summarized in Extended Data Table 1. Model quality was further assessed using MolProbity^81^ and Ramachandran analysis (Extended Data Table 1). Representative densities are shown in Extended Data Fig. 3. All structural figures were prepared in ChimeraX^77^.

### Protein sequence alignment

Multiple sequence alignments in Figs. 3d (top and middle panels), 4f, and Extended Data Fig. 8 were obtained using Clustal Omega^82,83^ (https://www.ebi.ac.uk/jdispatcher/msa/clustalo) and rendered with ESPript 3.0^84^.

For structure-based sequence alignment in Fig. 3d (bottom panel) and Extended Data Fig. 6d, the structures were superimposed using ChimeraX, yielding the aligned protein sequences annotated via ESPript 3.0 (https://espript.ibcp.fr/ESPript/ESPript/).

### Splicing analysis

Spicing of RBM5-dependent cassette exons was assayed by RT-PCR in RBM5-null cells, RBM10-mutant cells (RBM5^−/−^; RBM10^−/CE^), and 293Flp-In cells expressing RBM5-Flag, as described^27^.

Total RNA was extracted from pelleted cells using TRIzol (Thermo Fisher Scientific). DNA was removed with TURBO DNase (Thermo Fisher Scientific). Reverse transcription was performed with SuperScript IV (Thermo Fisher Scientific) using d(T)20 reverse primer cDNAs were amplified for 15-20 PCR cycles with Phusion Hot Start II DNA Polymerase (Thermo Fisher Scientific) and 5’-radiolabeled primers, resolved by denaturing PAGE, and detected by phosphorimaging on an Amersham Typhoon instrument. The resulting bands are quantified using ImageQuant (GE Healthcare Bio-Sciences).

## Acknowledgments

This project is supported by NIH grants R01GM071940 to Z.H.Z., R35 GM136426 to D.L.B. and R01 GM127473 to A.D. We acknowledge the use of resources at the Electron Imaging Center for Nanosystems, supported by UCLA, and grants from the NIH (S10RR023057, S10OD018111 and UL1TR001881) and NSF (DBI-1338135 and DMR-1548924).

## Author contributions

Z.H.Z., D.L.B., S.L. and A.D. conceived the project and supervised the research. S.L. and A.D. designed the experimental protocols. A.D, J. H. and C.-H.L. developed the isolation procedure, prepared the EM samples and performed the splicing assays. S.L. prepared the EM grids and collected cryo-EM images. S.L. and T.S. determined the 3D structures and built atomic models. S.L., A.D. and T.S. made figures. S.L., A.D., D.B. and Z.H.Z. interpreted the results and wrote the original manuscript. S.L. completed manuscript writing, figure preparation and final interpretation after relocating to Shandong University; no NIH funds or research resources were used for these activities. All authors contributed to the editing of the manuscript.

## Competing interests

We declare no competing interests.

## Data availability

The cryo-EM maps and atomic coordinates have been deposited to the EMDB (EMD-74089,74082,74090,74099,74100) and PDB (9ZE2,9ZE0,9ZE3,9ZEC,9ZED) databases, respectively. All other data are available within the main text or the Extended Data. Raw data and source images are available in the Supplementary Information. Source data are provided with this paper.

## Supplementary Materials

Reporting Summary

## Extended Data Figures

**Extended Data Fig. 1.**
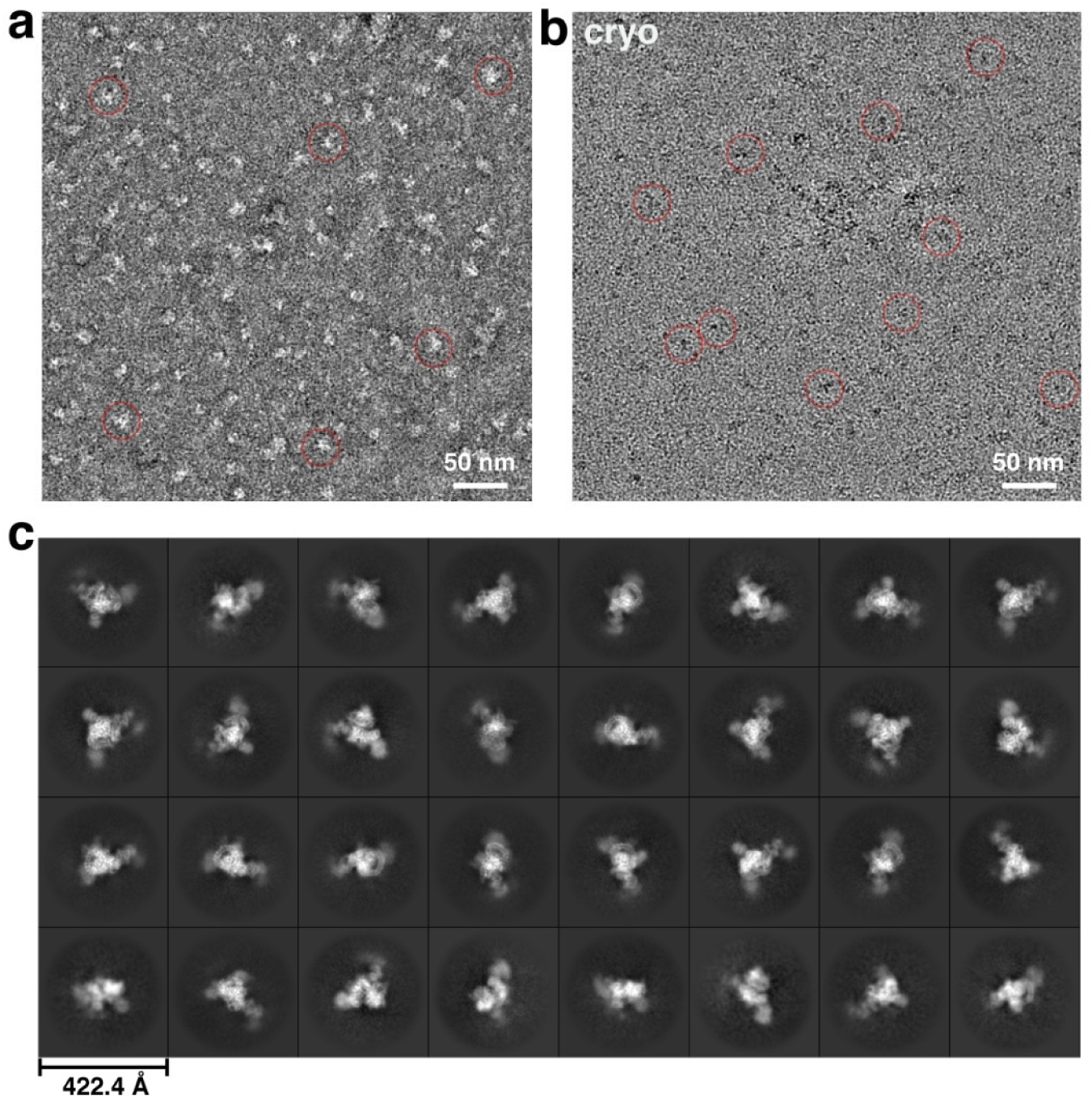
Representative cryo-EM images and 2D analyses. **a-b,** Negative staining (**a**) and drift-corrected cryo-EM (**b**) micrographs of native RNP from chromatin-associated HMW nuclear material. Representative particles are shown in red circles. **c**, Representative 2D class averages of the chromatin-derived RNP.

**Extended Data Fig. 2.**
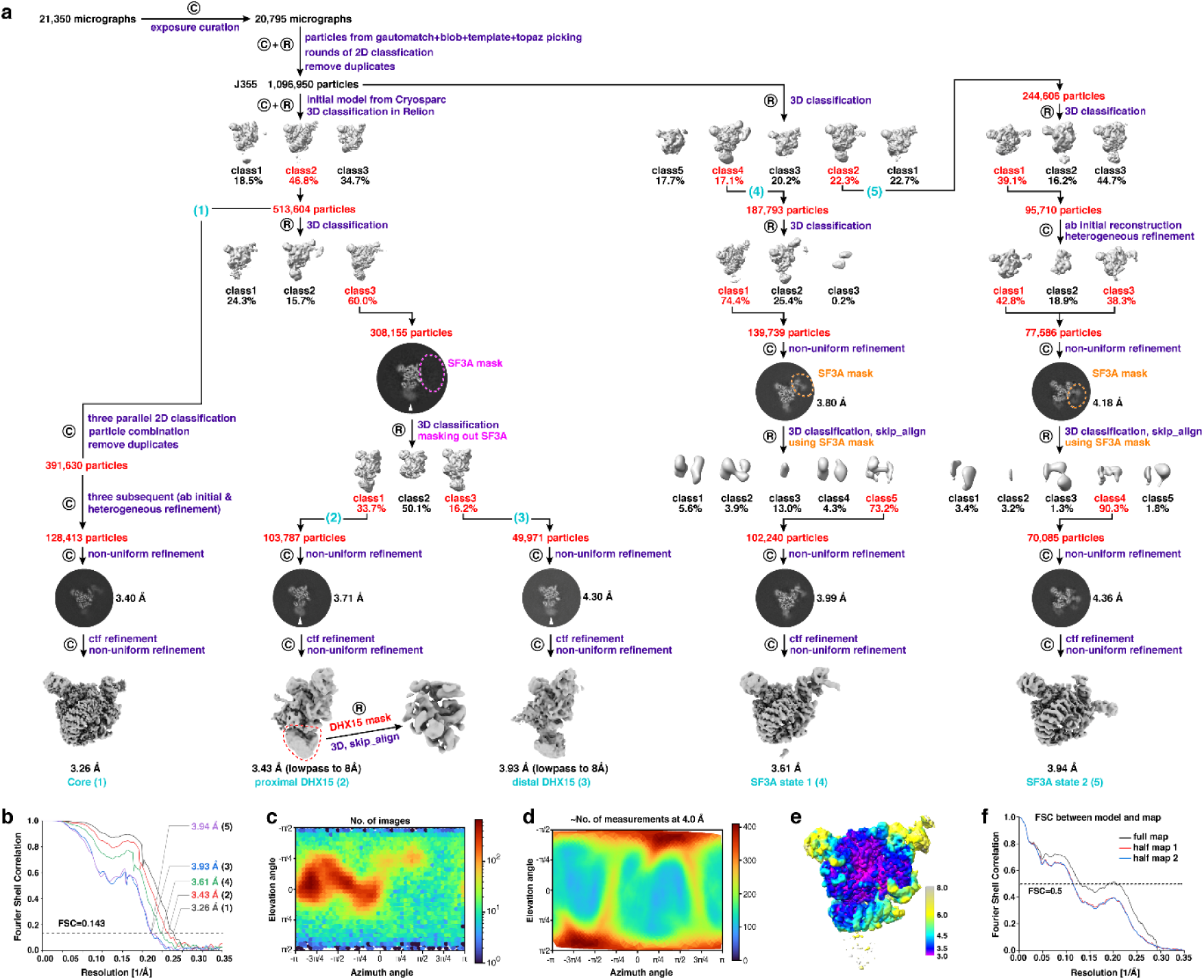
Cryo-EM structural determination of the in vivo purified spliceosome. **a**, Data processing workflow. The mask used in focused 3D classification are outlined with colored dashed lines. Data processing in RELION and cryoSPARC is denoted by a circular inscribed “R” and “C”, respectively. Steps used for processing the different states of the chromatin-derived RNP are indicated by the numbered labels shown in parentheses. See Methods for more details. **b**, FSC as a function of spatial frequency demonstrating the resolution of chromatin-derived RNPs. **c-d**, View direction distribution histogram (**c**) and posterior precision plot (**d**) show view diversity of all particles used for the final map of the RNP core (from cryoSPARC). **e**, Local resolution estimation of the RNP core (from cryoSPARC) using a local FSC threshold of 0.5. **f**, FSC curves between the proximal-DHX15 model containing RBM5, DHX15 and U2SURP and the corresponding cryo-EM maps. The generally similar appearances between the FSC curves obtained with half maps with (red) and without (blue) model refinement indicate that the refinement of the atomic coordinates did not suffer from severe over-fitting.

**Extended Data Fig. 3.**
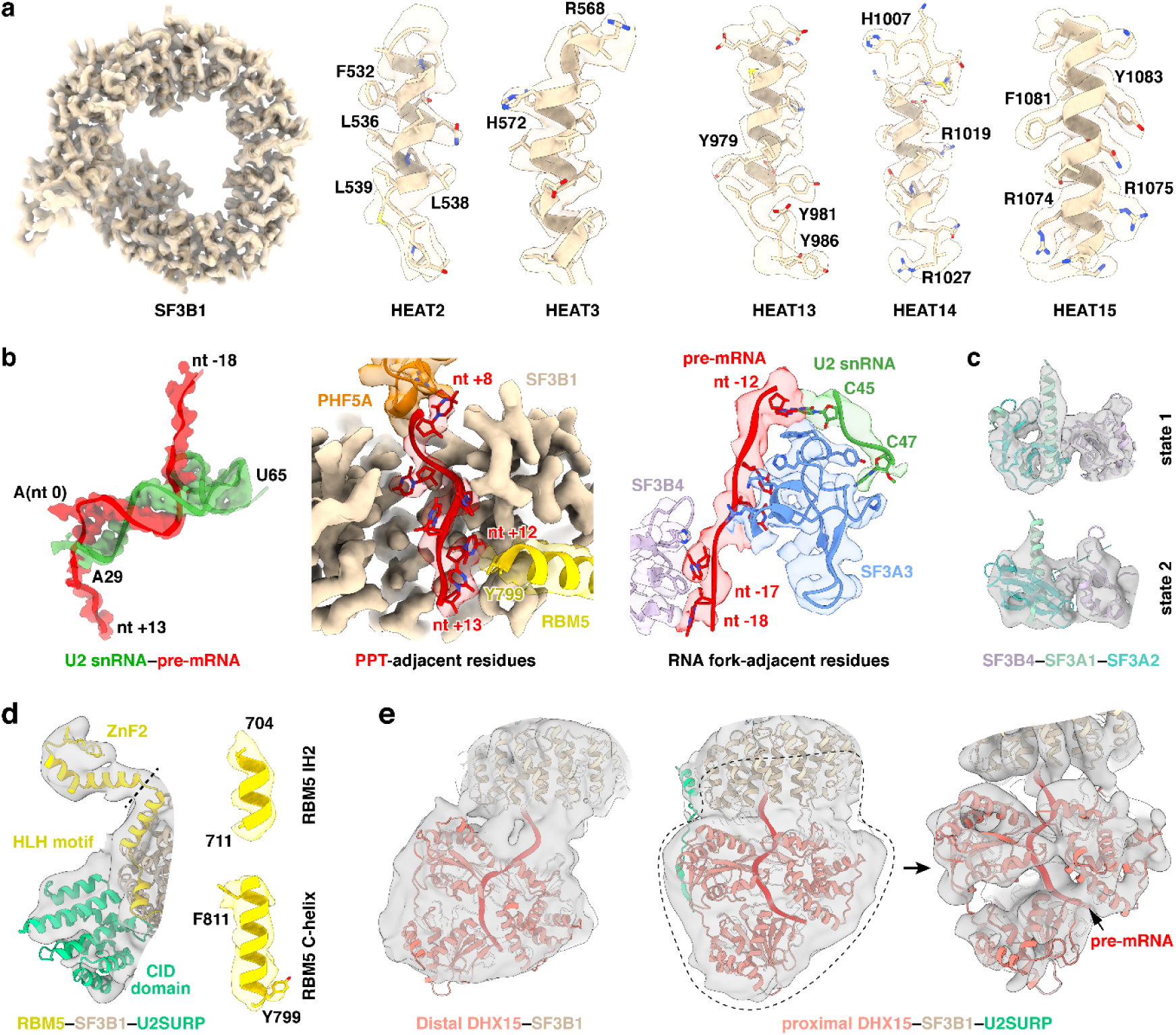
Representative cryo-EM density maps of the chromatin-derived RNP. **a**, SF3B1 displayed using the 3.26-Å sharpened map (core state). **b**, RNA elements and surrounding residues shown using the 3.43-Å sharpened map (proximal DHX15 state). **c**, Two conformations of the SF3B4–SF3A1–SF3A2 module shown using Gaussian low-pass–filtered maps of the 3.61-Å (SF3A state 1) and 3.94-Å unsharpened maps (SF3A state 2), respectively. **d**, Left: RBM5–SF3B1–U2SURP displayed using a Gaussian low-pass–filtered version of the 3.43-Å unsharpened map (proximal DHX15 state); right: RBM5 helices shown using the 3.43-Å sharpened map (proximal DHX15 state). **e**, Left: DHX15 shown using a Gaussian low-pass–filtered 3.43-Å unsharpened map (proximal DHX15 state); middle and right: DHX15 shown using Gaussian low-pass–filtered 3.93-Å unsharpened maps (distal DHX15 state).

**Extended Data Fig. 4.**
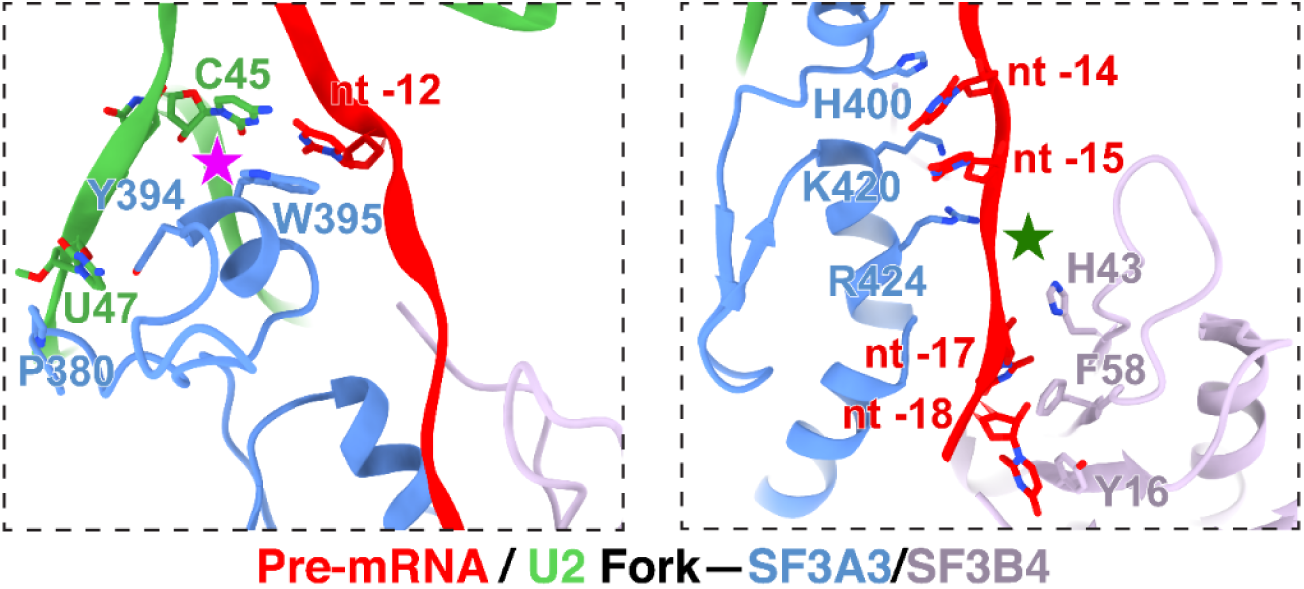
Residues engaging the forked U2 snRNA–pre-mRNA duplex upstream of BP.

**Extended Data Fig. 5.**
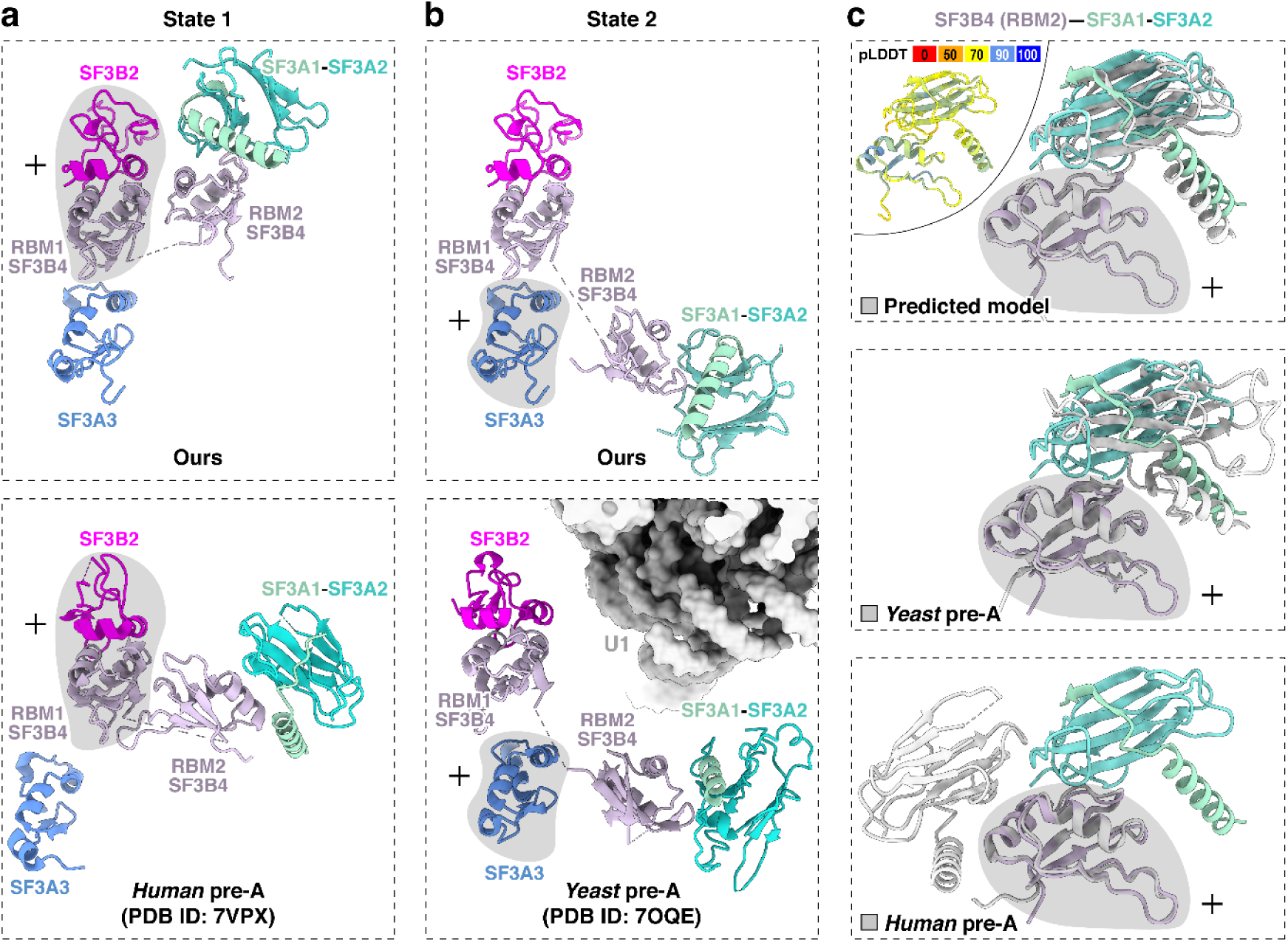
Structure of SF3B4–SF3A2–SF3A1 in two distinct states. Grey shading indicates regions used for structural superposition. **a,b,** Comparison of our SF3B4–SF3A2–SF3A1 (top) with *human* and *yeast* pre-A structures (bottom). State 1 aligns more closely with the *human* pre-A conformation (a), whereas state 2 resembles the *yeast* pre-A conformation (b). **c,** Comparison of our SF3B4 RRM2–SF3A1–SF3A2 interface (color-coded) with the AlphaFold-predicted, *yeast* pre-A, and *human* pre-A structures (grey). Top left: The predicted structure is colored by pLDDT scores, indicating the confidence level of each region.

**Extended Data Fig. 6.**
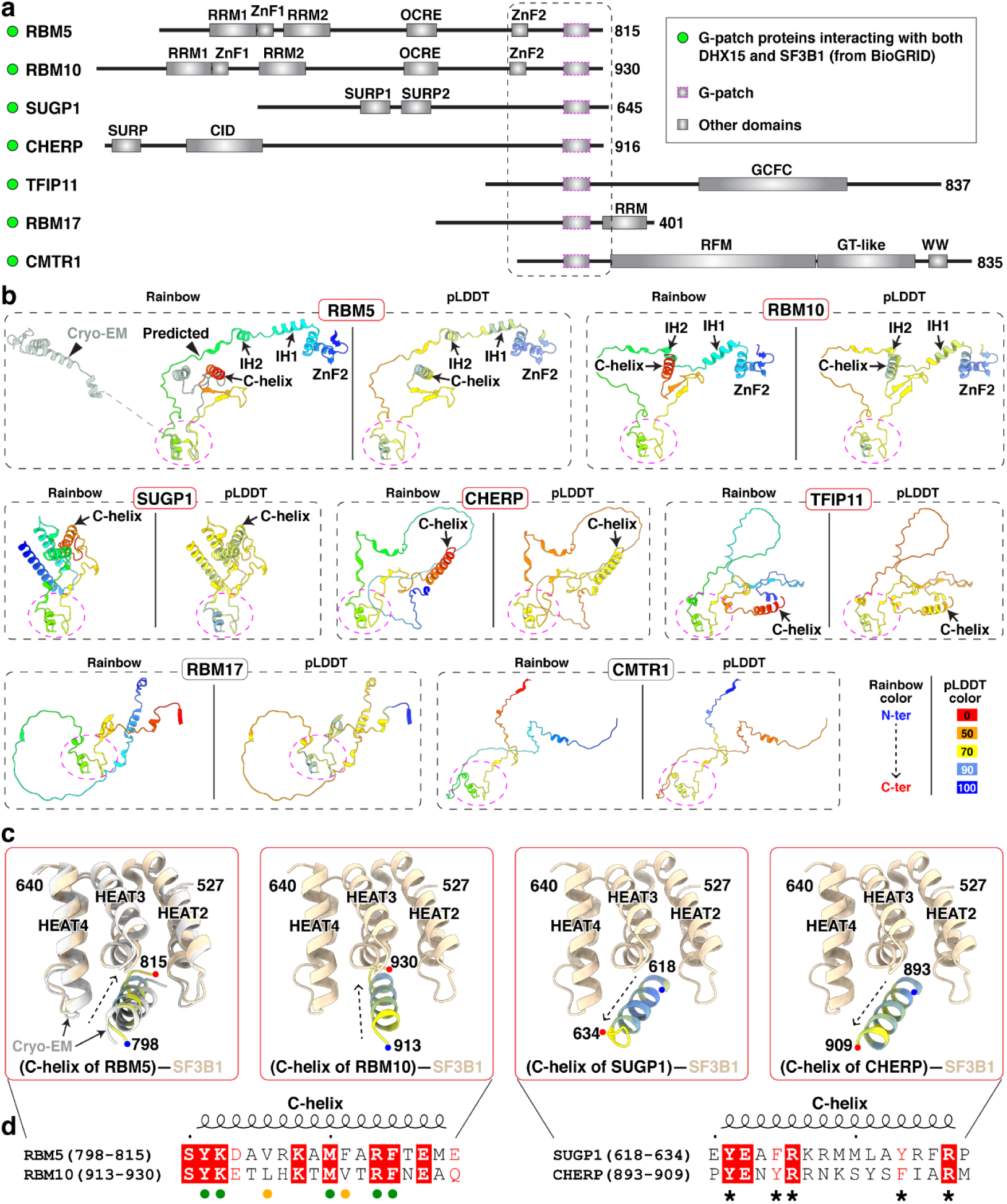
Putative SF3B1 binding by DHX15-associated G-patch proteins. **a,** Domain organization of BioGRID-identified G-patch proteins interacting with DHX15 and SF3B1. Proteins are aligned at the G-patch domain (purple dashed box), with adjacent N- and C-terminal regions boxed. Boundary of the dash box is defined by the resolved RBM5 structure. **b,** AlphaFold 3 structure prediction of G-patch-containing fragments, corresponding to the boxed regions in **(a)**. The G-patch is consistently oriented across all predicted structures (magenta circle). The left panels of all targets are shown in a rainbow color scheme from N to C terminus, while the right panels are colored by pLDDT. Regions with pLDDT > 90 are expected to be modelled to high accuracy; regions with pLDDT between 70 and 90 are expected to be modelled well; regions with pLDDT between 50 and 70 are modelled with low confidence and should be treated with caution. **c,** AlphaFold 3 predictions of G-patch protein C-helix interactions with SF3B1. All C-helices are colored by pLDDT, with our cryo-EM structure in grey (left panel) and SF3B1 in light tan. **d,** Sequence alignment of the C-helix with a conserved orientation when bound to SF3B1. Green and yellow circles mark identical residues in both RBM5 and RBM10 and those with conserved hydrophobicity, respectively. Asterisks denote conserved residues between SUGP1 and CHERP that are predicted to contribute to SF3B1 binding.

**Extended Data Fig. 7.**
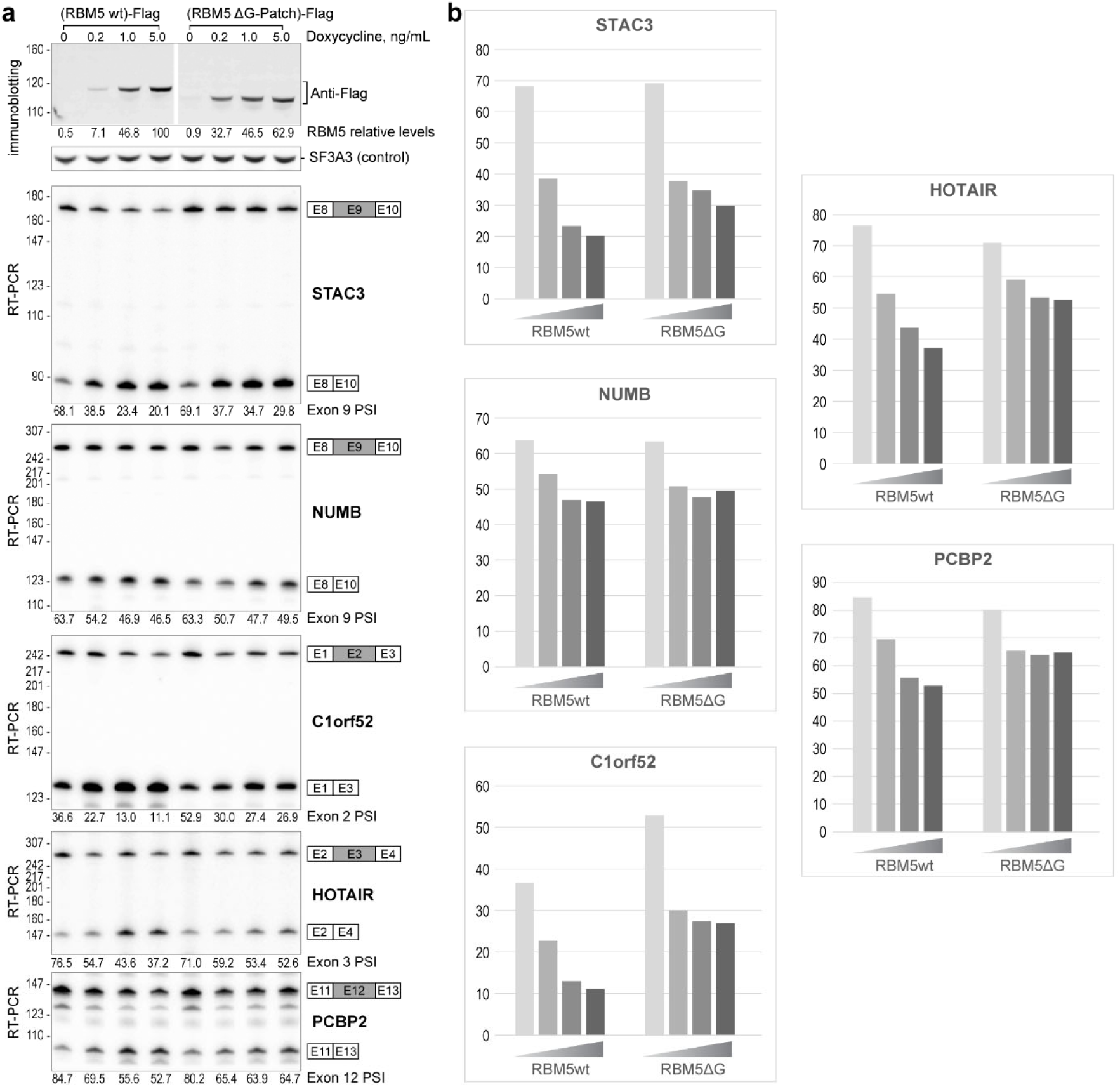
Regulatory effect of the RBM5 G-patch domain on splicing of exons in STAC3, NUMB, C1orf52, HOTAIR, and PCBP2. **a,** Top: Expression levels of Flag-tagged RBM5 and RBM5 ΔG-patch, quantified by immunostaining, with SF3A3 as a normalization control. Bottom: RT-PCR analysis of endogenous pre-mRNA splicing using radiolabeled flanking primers, followed by denaturing PAGE and phosphorimaging. Quantified exon inclusion levels are displayed below each RT-PCR result. **b,** Graph of quantified exon inclusion levels.

**Extended Data Fig. 8.**
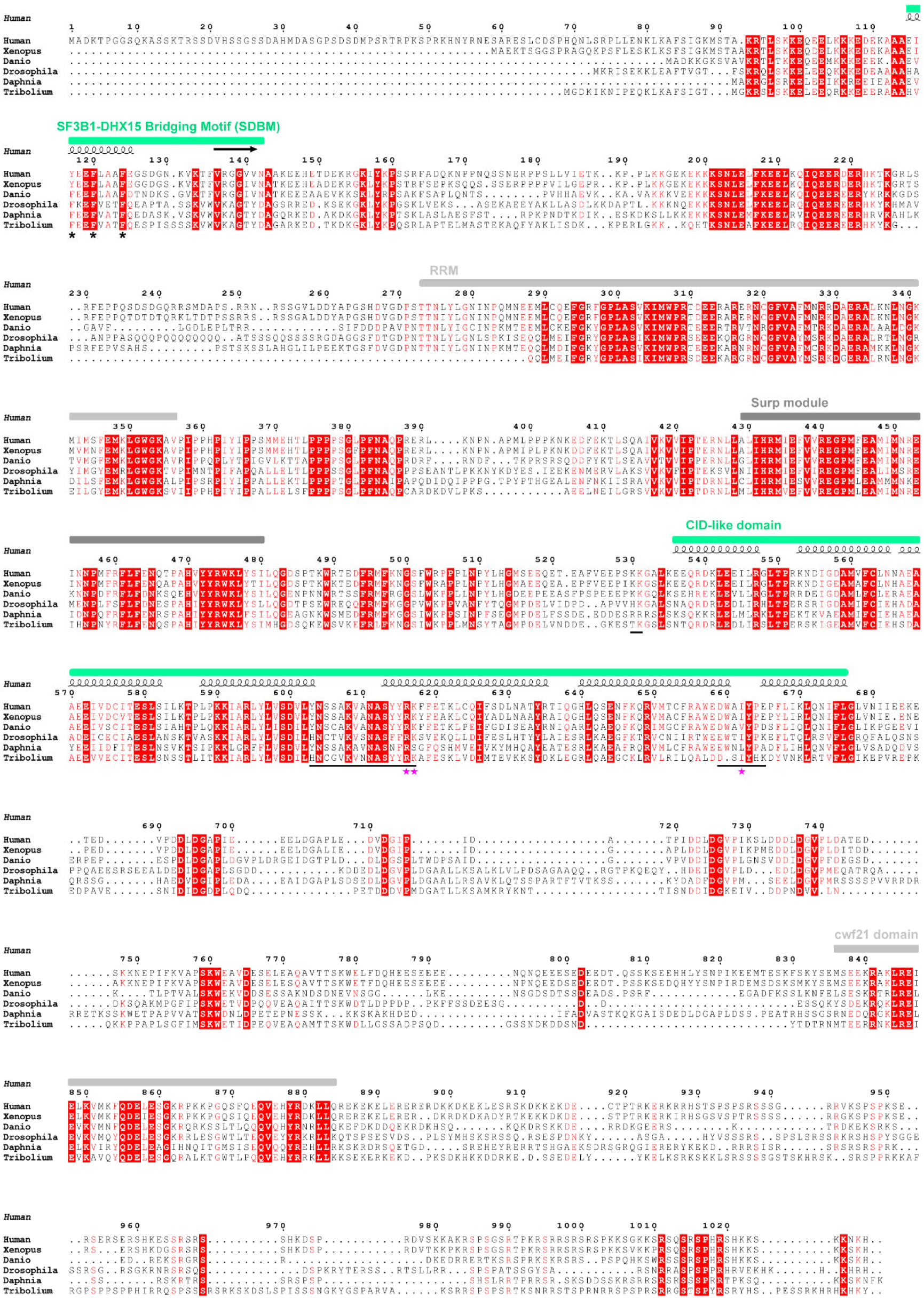
Multiple sequence alignment of the U2SURP proteins from different species. Domain organization of the full sequence and secondary structure of modeled regions are annotated at the top. Residues or regions mediating side-chain interactions are marked with asterisks and underlines, with single missense variants from COSMIC cancer database indicated by magenta stars at the bottom.

**Extended Data Fig. 9.**
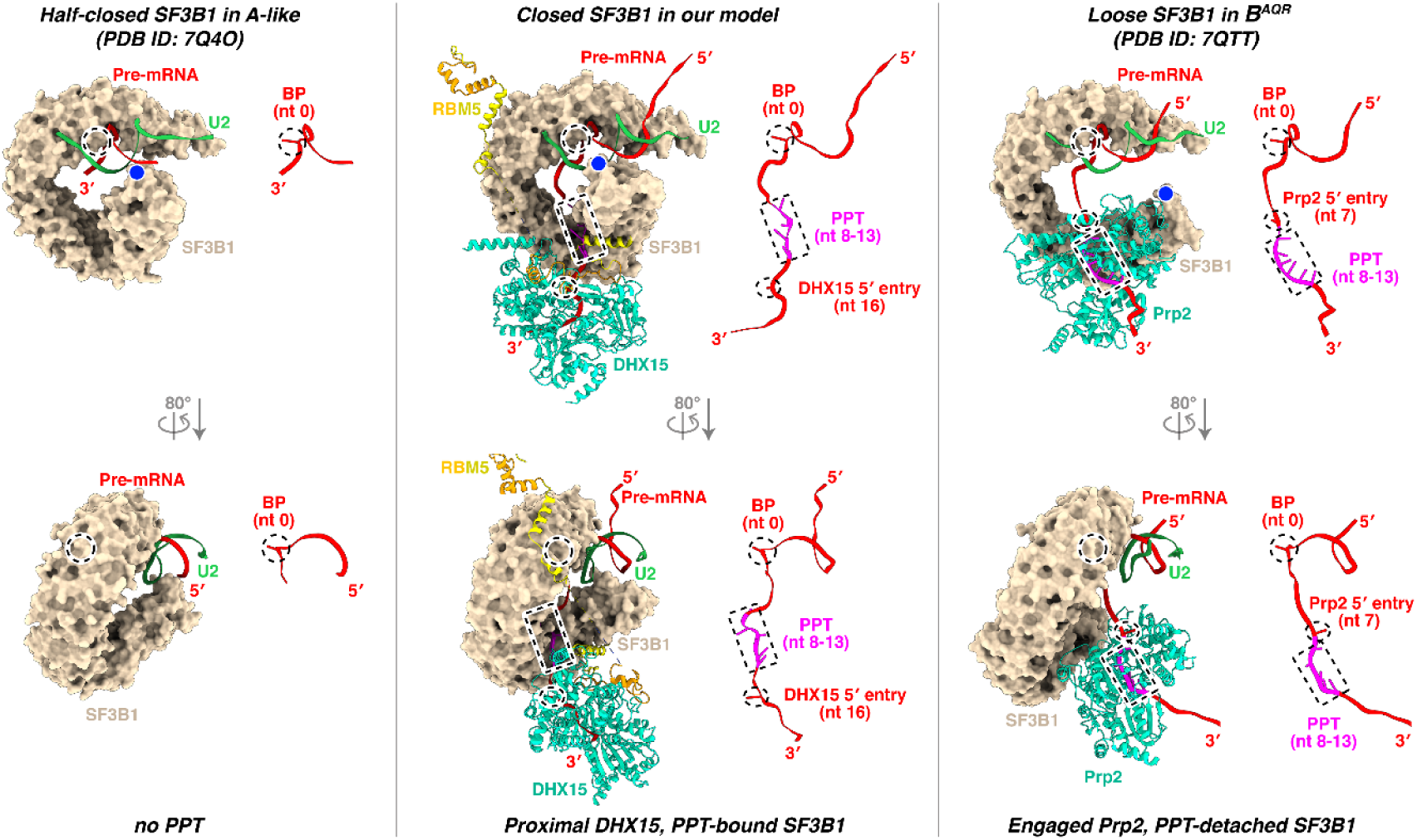
Structural transitions of DHX15 and pre-mRNA from the closed state of SF3B1 to its half-closed and loose states. For clarity, only the BS–U2 duplex with flanking nucleotides, SF3B1, and the DExH helicases (DHX15 in our model and Prp2 in B^AQR^) are shown, with RBM5 included in our model. Nucleotides corresponding to the BP, the likely PPT, and the helicase 5′-entry are marked by dashed circle, rectangle, and ellipse, respectively, and displayed as sticks in the right panels when present. Structures were superimposed using the C-terminal HEAT repeats of SF3B1, with conformational changes assessed by the relative displacement between the BP (dashed circle) and the N-terminal HEAT repeat (blue dot) in the left panels.

## Notes

### Competing Interest Statement

The authors have declared no competing interest.

