## Supplemental Table 1 for "The tumour suppressor RBM5 activates the helicase DHX15 to regulate splicing"

**Extended Data Table 1. Cryo-EM data collection, refinement and validation statistics**

| Structural states | Core | Proximal DHX15 | Distal DHX15 | SF3A state 1 | SF3A state 2 |
| --- | --- | --- | --- | --- | --- |
| EMDB | EMD-74089 | EMD-74082 | EMD-74090 | EMD-74099 | EMD-74100 |
| PDB | 9ZE2 | 9ZE0 | 9ZE3 | 9ZEC | 9ZED |
| Data collection and processing |  | | | | |
| Microscope | Titan Krios G1 | | | | |
| Voltage (kV) | 300 | | | | |
| Camera | K3 | | | | |
| Magnification | 81K | | | | |
| imaging mode | super-resolution | | | | |
| Pixel size (Å) | 0.55 | | | | |
| Electron exposure (e^-^/Å^2^) | ~50 | | | | |
| Defocus range (μm) | -1.5 to -3.0 | | | | |
| Symmetry imposed | C1 | | | | |
| Micrographs processed | 20,795 | | | | |
| Final particles | 128,413 | 103,787 | 49,971 | 102,240 | 70,085 |
| Map resolution | 3.26 | 3.43 | 3.93 | 3.61 | 3.94 |
| FSC threshold | 0.143 | 0.143 | 0.143 | 0.143 | 0.143 |
| Model refinement and validation |  | | | | |
| Map sharpening B-factor (Å^2^) | -127.6 | -119.8 | -119.5 | -127.1 | -116.4 |
| Model resolution (Å) | 2.21 | 2.95 | 6.5 | 3.4 | 4.1 |
| FSC threshold | 0.5 | 0.5 | 0.5 | 0.5 | 0.5 |
| Model composition |  | | | | |
| Non-hydrogen atoms | 24,526 | 38,259 | 37,998 | 27,423 | 26,632 |
| Protein residues | 2,912 | 4,571 | 4,530 | 3,264 | 3,167 |
| Nucleotide residues | 69 | 79 | 81 | 69 | 68 |
| R.m.s. deviations |  | | | | |
| Bonds length (Å) | 0.002 | 0.003 | 0.002 | 0.002 | 0.002 |
| Bonds angles (Å) | 0.522 | 0.492 | 0.498 | 0.477 | 0.495 |
| **Validation** |  | | | | |
| MolProbity score | 1.46 | 1.48 | 1.49 | 1.40 | 1.51 |
| Clashscore | 4.10 | 5.28 | 5.19 | 4.46 | 5.88 |
| Poor rotamers (%) | 0.12 | 0.05 | 0.03 | 0.25 | 0.07 |
| Ramachandran plot |  | | | | |
| Favored (%) | 0 | 0 | 0 | 0 | 0 |
| Allowed (%) | 3.85 | 3.23 | 3.34 | 3.07 | 3.13 |
| Disallowed (%) | 96.15 | 96.77 | 96.66 | 96.93 | 96.87 |
